# A cellular census of healthy lung and asthmatic airway wall identifies novel cell states in health and disease

**DOI:** 10.1101/527408

**Authors:** F.A. Vieira Braga, G. Kar, M. Berg, O.A. Carpaij, K. Polanski, L.M. Simon, S. Brouwer, T. Gomes, L. Hesse, J. Jiang, E.S. Fasouli, M. Efremova, R. Vento-Tormo, K. Affleck, S. Palit, P. Strzelecka, H.V. Firth, K.T.A. Mahbubani, A. Cvejic, K.B. Meyer, K. Saeb-Parsy, M. Luinge, C.-A. Brandsma, W. Timens, I. Angelidis, M. Strunz, G.H. Koppelman, A.J. van Oosterhout, H.B. Schiller, F.J. Theis, M. van den Berge, M.C. Nawijn, S.A. Teichmann

## Abstract

Human lungs enable efficient gas exchange, and form an interface with the environment which depends on mucosal immunity for protection against infectious agents. Tightly controlled interactions between structural and immune cells are required to maintain lung homeostasis. Here, we use single cell transcriptomics to chart the cellular landscape of upper and lower airways and lung parenchyma in health. We report location-dependent airway epithelial cell states, and a novel subset of tissue-resident memory T cells. In lower airways of asthma patients, mucous cell hyperplasia is shown to stem from a novel mucous ciliated cell state, as well as goblet cell hyperplasia. We report presence of pathogenic effector Th2 cells in asthma, and find evidence for type-2 cytokines in maintaining the altered epithelial cell states. Unbiased analysis of cell-cell interactions identify a shift from airway structural cell communication in health to a Th2-dominated interactome in asthma.

## Introduction

The lung plays a critical role in both gas exchange and mucosal immunity, and its anatomy serves these functions through (1) the airways that lead air to the respiratory unit, provide mucociliary clearance, and form a barrier against inhaled particles and pathogens; and (2) the alveoli, distal saccular structures where gas exchange occurs. Acute and chronic disorders of the lung are a major cause of morbidity and mortality worldwide^1^. To better understand pathogenesis of lung disease, it is imperative to characterise the cell types of the lung and understand their interactions in health^2,3^ and disease. The recent identification of the ionocyte as a novel airway epithelial cell-type^4,5^ underscores our incomplete understanding of the cellular landscape of the lung, which limits our insight into the mechanisms of respiratory disease, and hence our ability to design therapies for most lung disorders.

We set out to profile lung-resident structural and inflammatory cells and their interactions by analysing healthy human respiratory tissue from four sources: nasal brushes, endobronchial biopsies and brushes from living donors, and tissue samples from lung resections and transplant donor lungs. Our single cell analysis identifies differences in the proportions and transcriptional phenotype of structural and inflammatory cells between upper and lower airways and lung parenchyma. Using an unbiased approach to identify tissue-resident CD4 T cells in airway wall, we identify a novel tissue migratory CD4 T cell (TMC) that harbours features of both circulating memory cells and of tissue resident memory cells (TRM) CD4 T cells. We demonstrate that many disease-associated genes have highly cell type-specific expression patterns. This holds true for both rare disease-associated genes, such as CFTR mutated in cystic fibrosis, as well as genes associated with a common disease such as asthma.

In addition, we evaluate the altered cellular landscape of the airway wall in chronic inflammatory disease using bronchial biopsies from asthma patients. We identify a novel epithelial cell state highly enriched in asthma, the mucous ciliated cell. Mucous ciliated cells represent a transitioning state of ciliated cells with molecular features of mucus production, and contribute to mucous cell hyperplasia in this chronic disease. Other changes associated with asthma include increased numbers of goblet cells, intraepithelial mast cells and pathogenic effector Th2 cells in airway wall tissue. We examine intercellular communications occurring in the healthy and asthmatic airway wall, and reveal a remarkable loss of epithelial communication and a concomitant increase in Th2 cell interactions. The newly identified TMC subset interacts with epithelial cells, fibroblasts and airway smooth muscle cells in asthma. Collectively, these data generate novel insights into epithelial cell changes and altered communication patterns between immune and structural cells of the airways, that underlie asthmatic airway inflammation.

### A human lung cell census identifies macro-anatomical patterns of epithelial cell states across the human the respiratory tree

The cellular landscape along the 23 generations of the airways in human lung is expected to differ both in terms of relative frequencies of cell types and their molecular phenotype^6^. We used 10x Genomics Chromium droplet single-cell RNA sequencing (scRNA-Seq) to profile a total of 36,931 single cells from upper and lower airways, and lung parenchyma (Figure 1A, B). We profiled nasal brushes, and (bronchoscopic) brushes and biopsies from airway wall (third to sixth generation) from healthy volunteers. For parenchyma (small respiratory airways and alveoli), we obtained lung tissue from deceased transplant donors, also analysed on the 10x platform, and from non-tumour resection tissue from lung cancer patients, analysed on a bespoke droplet microfluidics platform based on the Dropseq protocol ^7^.

**Figure 1.**
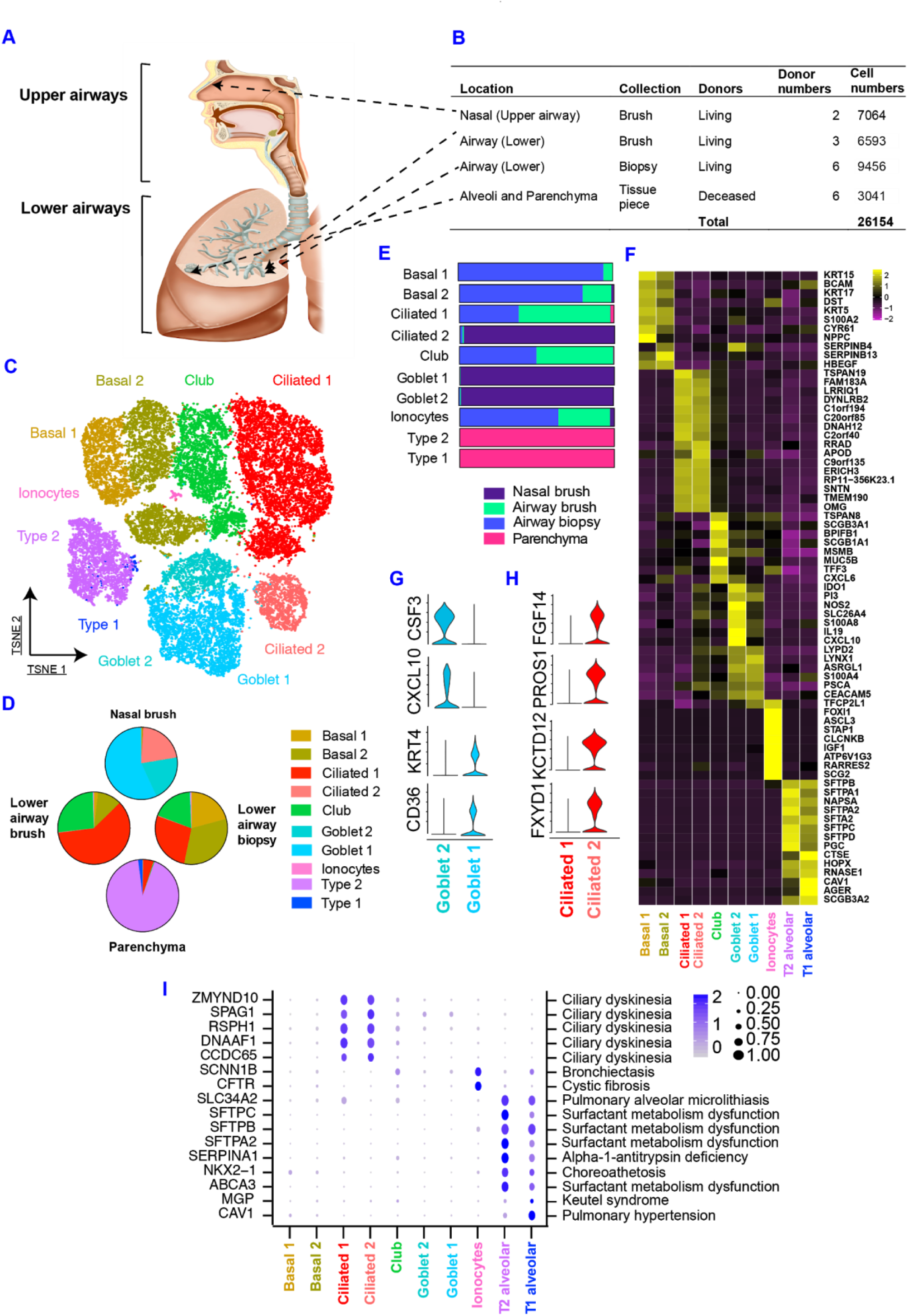
A human lung cell census identifies zonation of novel epithelial cell states across macro-anatomical location. **(A)** Schematic illustration depicting anatomical regions analysed in this manuscript. **(B)** Table with the details of anatomical region, tissue source, donors and cell numbers present in this figure. **(C)** tSNE displaying the major epithelial clusters present in the full extent of the human respiratory tree. **(D**) Pie charts depicting the cellular composition by anatomical region. **(E)** Horizontal slice bar depicting the anatomical distribution of each cell type identified **(F)** Heatmap depicting the average expression levels per cluster of the top differentially expressed markers in each cluster. **(G)** Violin plots of selected markers identified by differential expression analysis comparing the two goblet subsets to each other. **(H)** Violin plots of selected markers identified by differential expression analysis of ciliated 1 versus ciliated 2 clusters. **(I)** Dot plot depicting gene expression levels and percentage of cells expressing genes associated with specific lung phenotypes according to the Online Mendelian Inheritance in Man (OMIM) database. Only genes present in the top 50 (per cluster) of our list of differentially expressed genes are depicted in (I). All the differential expression analysis were performed using Wilcoxon rank sum test in Seurat^48^.

Integration of the data from nasal epithelium, airway wall and parenchymal tissue reveals a diversity of epithelial, endothelial, stromal and immune cells, with approximately 21 coarse-grained cell types in total (Figures 1 and 2, Extended Figure 1), that can be explored in user-friendly web portal (www.lungcellatlas.org). Analysis of parenchymal lung tissue from resection material using Dropseq led to the identification of 15 coarse-grained cell populations (epithelial and non-epithelial) (Extended Figure 2). Using MatchSCore^8^ to quantify the overlap between cell type marker signatures between the two datasets revealed an extensive degree of overlap in cell type identities (Extended Figure 2). In our analysis below, we first concentrate on epithelial cells (Figure 1), and then focus on the stromal and immune compartments (Figure 2).

**Figure 2.**
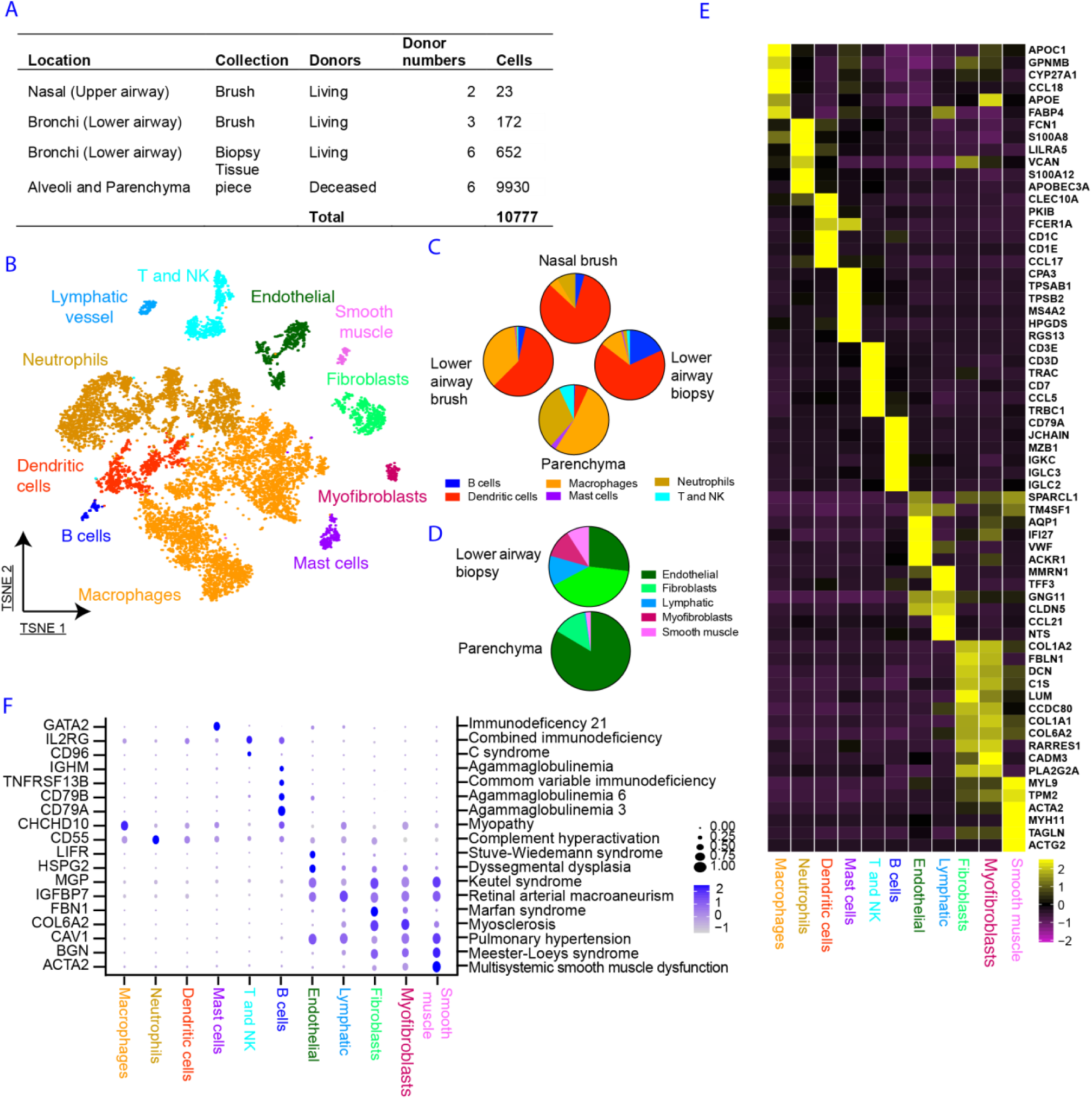
A cellular and molecular map of the stromal and immune components of across the upper and lower human respiratory airways. **(A)** Table with details of anatomical region, tissue source, donors and cell numbers present in this figure. **(B)** tSNE displaying the major immune and mesenchymal clusters present in the full extent of the human respiratory tree. **(C)** Pie charts depicting the cellular composition of immune cells by anatomical region. **(D)** Pie charts depicting the cellular composition of stromal cells in lower airway biopsies and parenchyma tissue. **(E)** Heatmap depicting the average expression levels per cluster of the top differentially expressed markers in each cluster. **(F)** Dot plot depicting gene expression levels and percentage of cells expressing genes associated with lung phenotypes according to the Online Mendelian Inheritance in Man (OMIM) database. Only genes present in the top 50 (per cluster) of our list of differentially expressed genes are depicted in (F). All the differential expression analysis was performed using Wilcoxon rank sum test in Seurat.

In the epithelial lineage, we identified a total of at least 10 cell types across the upper and lower airways and lung parenchyma (Figure 1C, Extended Data Figure 1). We detected multiple basal, club, ciliated and goblet cell states, as well as type-1 (T1) and type-2 (T2) alveolar cells, and the recently described ionocyte^4,5^ (Extended Figure 3). Both goblet and ciliated cells were present in the nasal epithelium (Figure 1D). In the lower airways, we detected basal, club and ciliated cells as well as ionocytes, but only very small numbers of goblet cells. T1 and T2 cells were, as expected, only found in the lung parenchyma (Figure 1E).

**Figure 3.**
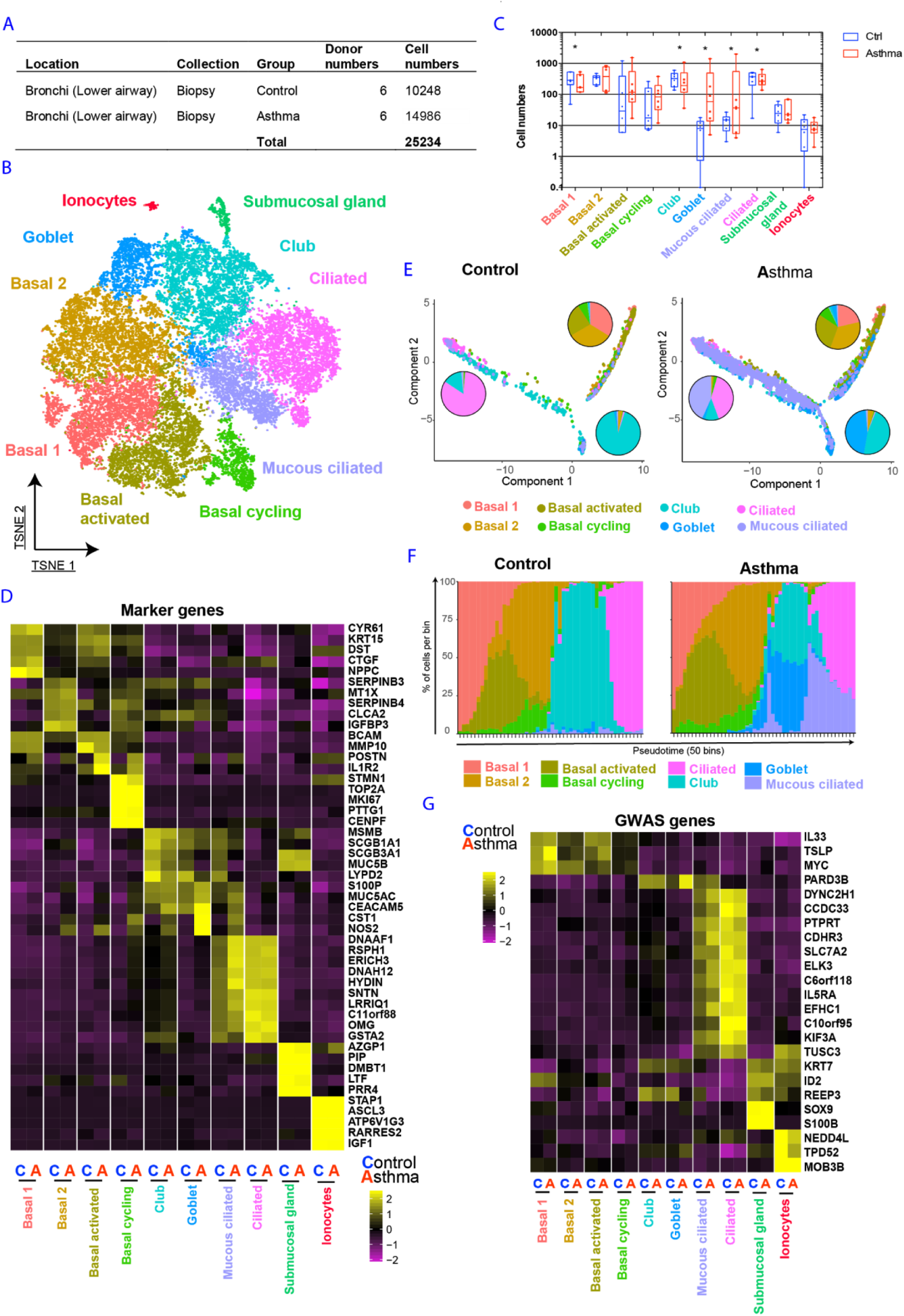
Distinct programs of epithelial cell differentiation in asthmatic *versus* healthy airways. **(A)** Table with overview and cell numbers for control and asthma volunteers analysed in this figure. **(B)** tSNE displaying all epithelial cells analysed coloured by their specific cluster assignment. **(C)** Box and whisker plots depicting cell numbers of control and asthma patients in each cluster**. (D)** Heatmap displaying the top five differentially expressed genes per cluster. **(E)** Pseudotime developmental trajectory analysis from Monocle2 depicting how each of the basal, secretory and ciliated subsets relate to each other. **(F)** Binned pseudotime analysis displaying how each subset is ordered in a one-dimensional continuous space. **(G)** Heatmap displaying the expression of asthma genes from GWAS. Only genes present in our list of differentially expressed genes are depicted for each cell cluster. Significance analysed using Fisher’s exact test corrected for multiple comparison using the Bonferroni method. Significance calculated using all the clusters present in figures 3 and 4, which were derived from the same set of samples. *represents p-value<0.001. n=6 controls and n=6 asthma. The differential expression analysis used for input in D and E was performed using Wilcoxon rank sum test in Seurat.

We did not identify specific clusters of tuft cells or neuroendocrine (NE) cells. Since cell types represented by a small fraction of the data might be missed by unsupervised clustering, we evaluated the expression of known marker genes for NE cells (CHGA, ASCL1, INSM1, HOXB5) and Tuft cells (DCLK1, ASCL2)^4^. NE marker genes identified a small number of cells, present only in lower airways, displaying a transcriptional profile consistent with that of NE cells (extended Figure 4). Tuft cell marker genes did not identify a unique cell population. Ionocytes were found in lower airways, and at very low frequency in upper airways, but were completely absent from the parenchyma. Comparison of the cell populations identified using the two different bronchoscopic sampling methods (brush versus biopsy) in lower airways showed that basal cells were captured most effectively in biopsies, while apical epithelial cells, such as ciliated and club cells were relatively overrepresented in the bronchial brushings (Figure 1D).

**Figure 4.**
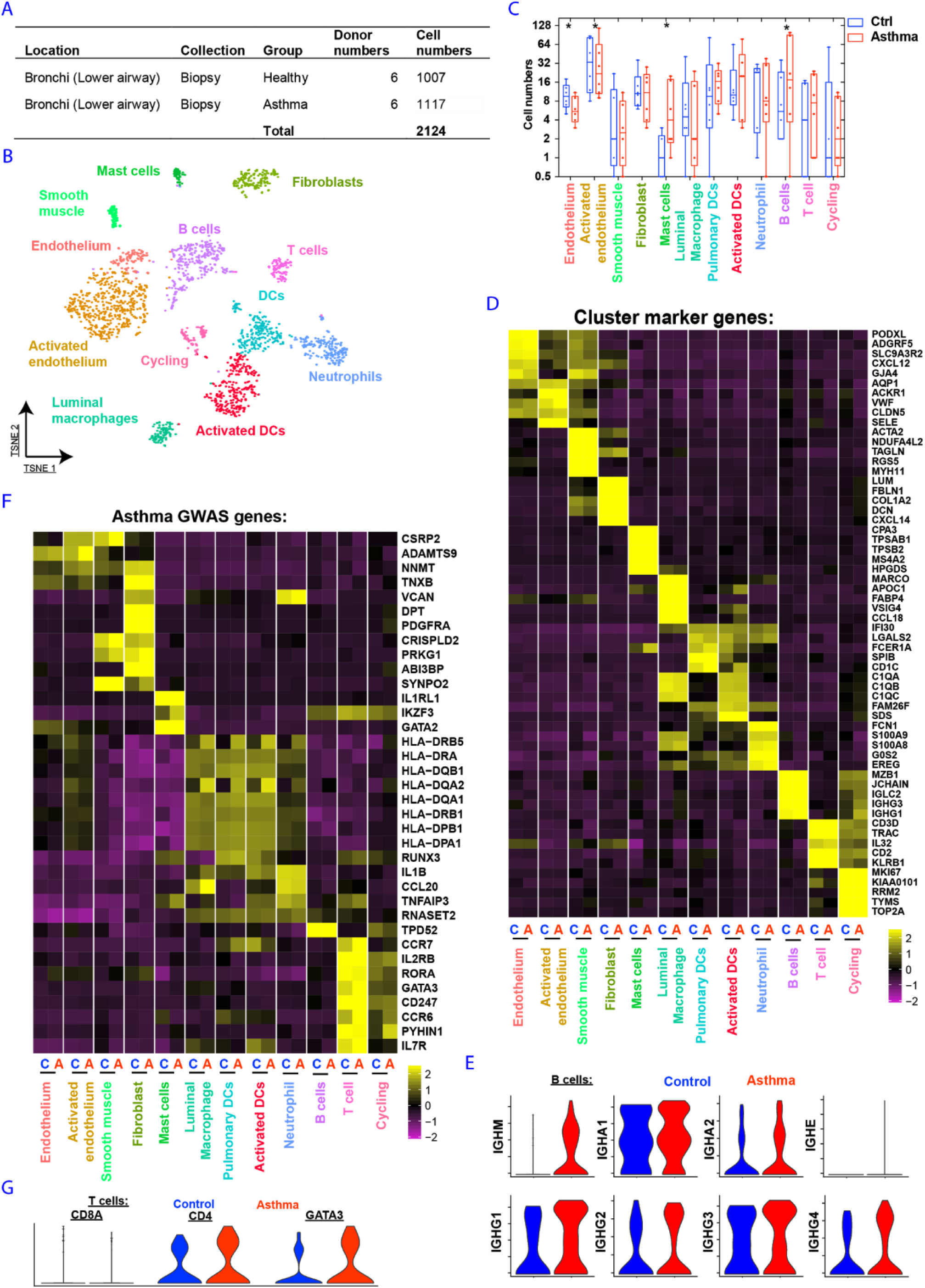
Remodelling of the stromal and Immune compartments in asthmatic airways. **(A)** Table with the number of donors and cells per volunteer group included in this figure. **(B)** tSNE depicting the immune and stromal cell types identified in the human airway combined dataset of healthy and asthmatic patients. **(C)** Box and whisker plots depicting the cell numbers of healthy and asthmatic cells in each cluster. **(D)** Heatmap displaying gene expression levels of the top 5 differentially expressed genes per cluster. **(E)** Violin plots depicting expression of immunoglobulin genes in B cells. **(F)** Heatmap displaying asthma GWAS gene expression per cluster. Only genes present in the top 50 (per cluster) of our list of differentially expressed genes are shown. **(G)** Violin plots of selected T cell markers in asthma patients. Significance calculated using all the clusters present in figures 3 and 4, which were derived from the same set of samples. *represents p-value<0.001. n=6 controls and n=6 asthma.

Our dataset allowed us to identify two discrete cell states in basal, goblet and ciliated epithelial cells. Some of these cell phenotypes were restricted to specific anatomical locations along the respiratory tract. Basal cells were present in both upper and lower airways, although at relatively low frequency in upper airways (Figure 1E). The two basal cell states corresponded to differentiation stages, with the less mature Basal 1 cell state expressing higher levels of *TP63* and *NPPC* in comparison to Basal 2 cells (Figure 1F and extended data 1), which were more abundant in bronchial brushes, suggesting a more apical localization for these more differentiated basal cells (Figure 1D). Goblet 1 and 2 cells were both characterized by high expression of CEACAM5, S100A4, MUC5AC and lack of MUC5B (Figure 1F and Extended Figures 1 and 4). Goblet 1 cells specifically express KRT4 and CD36 (Figure 1G and Extended Figure 4). Genes involved with immune function, such as *IDO1, NOS2, IL19, CSF3* (Granulocyte-colony stimulating factor) and *CXCL10 are* expressed at high levels in Goblet 2 cells (Figure 1G and Extended Figure 4). These molecules enriched in Goblet 2 cells are involved in recruitment of neutrophils, monocytes, dendritic cells and T cells^9^. Both goblet cells states are present in upper airway epithelium, with Goblet 1 cells being more frequent. In contrast, the Goblet 2 cell state was also present in lower airways, albeit at low abundance (Figure 1E).

Ciliated cell transcriptional phenotypes are also zonated in terms of their presence across macro-anatomical locations, with a discrete ciliated cell state more abundant in upper airways (Ciliated 2) compared to lower airways and parenchyma. Nasal epithelial Ciliated 2 cells express pro-inflammatory genes, such as *CCL20* (Extended data 3) and higher levels of metabolic genes (*ATP12A* and *COX7A1*) and vesicle transport (*AP2B1* and *SYT5*^*10*^) compared to the Ciliated 1 cell state. In contrast, the Ciliated 1 cells from lower airways specifically expressed genes involved in cytoprotection (*PROS1*^*11*^) and fluid reabsorption (*FXYD1*^*12*^) (Figure 1H and Extended Figure 4). Interestingly, comparison of the location-specific differences between ciliated and goblet cells identified a transcriptional signature specific for the upper airways present in both epithelial cell types (Extended Figure 4B).

Next, we assessed the contribution of specific epithelial cell types to Mendelian disease. Cell-type specific expression patterns of genes associated with Mendelian disorders (based on the Online Mendelian Inheritance in Man, OMIM database) confirm ionocytes as particularly high expressers of the *CFTR* gene, mutated in cystic fibrosis (Figure 1I). These cells also express *SCNN1B*, mutations of which can cause bronchiectasis, another feature of cystic fibrosis, suggesting a potential key pathological role for ionocytes in both bronchiectasis and cystic fibrosis. In addition, expression of *SERPINA1* (Figure 1I) was found to be enriched in type-2 alveolar epithelial cells, underscoring their role in alpha-1-antitrypsin deficiency^13^.

### Differential anatomical distribution of the stromal and immune components in the human respiratory tree

Next, we analysed the single cell transcriptomes of immune and stromal cells from the upper airways, lower airways and the lung parenchyma (Figure 2A). We identified immune clusters of myeloid (macrophages, neutrophils, dendritic cells (DCs) and mast cells) and lymphoid cells (T and NK cells, B cells; Figure 2B, and Extended Figure 5). Immune and stromal cell numbers and composition varied greatly across different anatomical regions (Figure 2A and 2C). Nasal brushes contained only a small number of immune cells, with the large majority being dendritic cells. In the lower airways, the fraction of inflammatory cells was significantly larger and relatively enriched for macrophages (Figure 2C and Extended Figure 5), which was directly confirmed by cell composition comparison of upper *versus* lower airway brushes obtained from the same donor (Extended Figure 5E).

**Figure 5.**
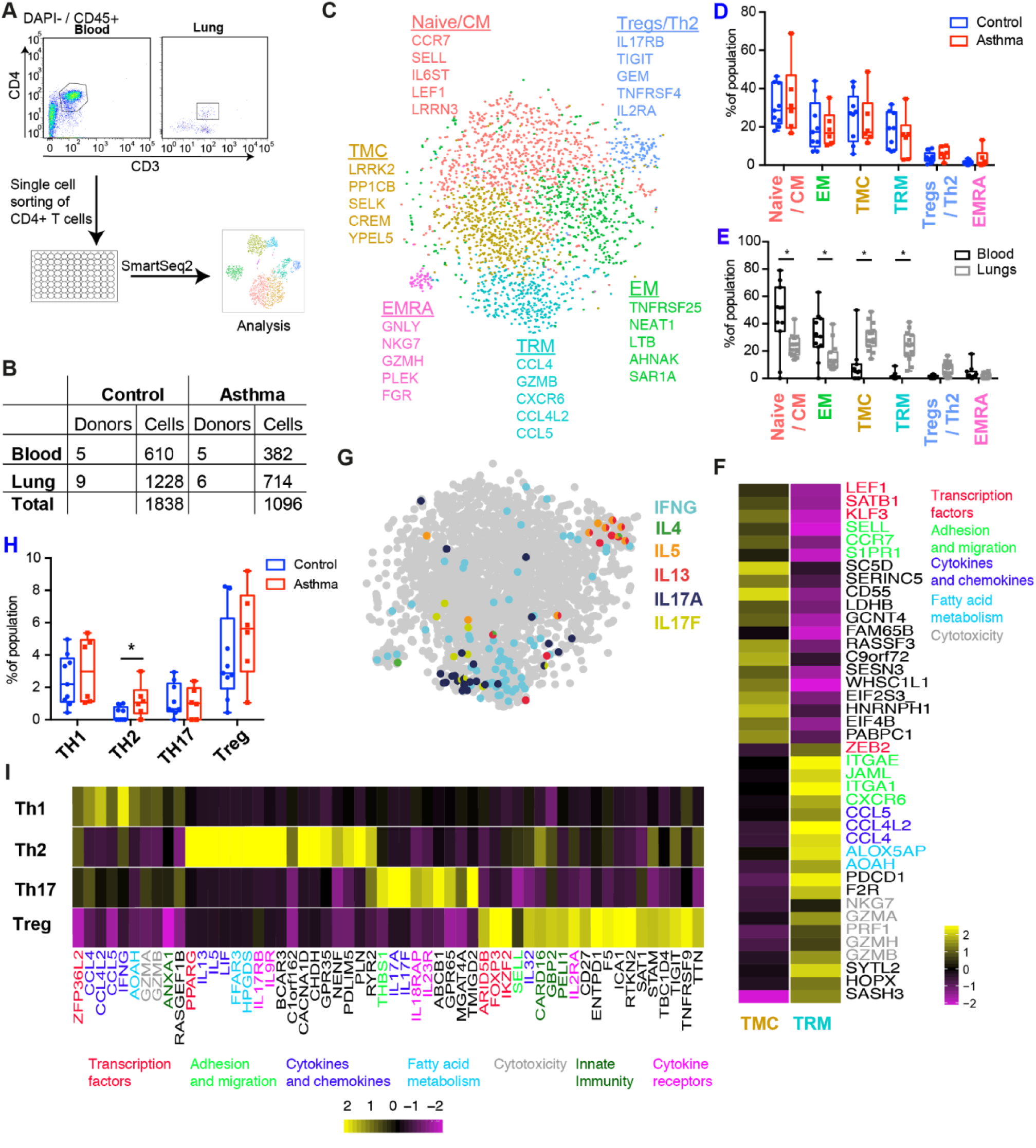
Pathogenic effector Th2 cells are enriched in asthmatic airways. **(A)** Schematic depicting experimental layout of single cell sorting of CD4 T cells from blood and lung airway biopsies. **(B)** Table with the number of donors by anatomical location for control and asthma groups. **(C)** tSNE displaying clusters of T cells identified by analysing the combined cells from blood and lung from control and asthma groups. **(D)** Box and whisker plots showing the cluster cell distributions from control and asthma patients. **(E)** Box and whisker plots depicting the cluster composition per donor according to the tissue source from which the cells were isolated. **(F)** Heatmap showing the average expression per cluster of genes differentially expressed between the two lung specific CD4 T cell populations. Gene names coloured according to functional categories. **(G)** tSNE depicting canonical cytokines from Th1, Th2 and Th17 cells. **(H)** Box and whisker plots showing the number of Th1, Th2 and Th17 cells defined by canonical cytokines expression and Tregs identified by unbiased clustering. **(I)** Heatmap of average cluster gene expression of markers differentially expressed between Th1, Th2, Th17 and Treg cells. Gene names coloured according to functional categories. Error bars in **(D), (E)** and **(H)** represent standard deviation. Significance analysed using Multiple t-tests corrected for multiple comparison using the Holm-Sidak method. * represents p-value<0.05. Patient numbers per tissue depicted in (B).

Macrophages show large donor variation in their phenotype (Extended figure 5), but they all share high expression of MARCO, CCL18 and genes involved in apolipoprotein metabolism (APOC1 and APOE) (Figure 2E). Lung neutrophils express high levels of the granulocyte markers S100A8, S100A12^14^ and LILRA5, a receptor poorly characterised in the lungs, that has been shown to have a proinflammatory function in synovial fluid macrophages ^15^ (Figure 2E). DCs were mostly myeloid, with high expression of CD1E, CD1C, CLEC10A (Figure 2E) and of FCER1A (IgE receptor) and CCL17, molecules known to play a key role in inflammatory conditions such as asthma^16^.

In the droplet RNAseq data sets, we could not distinguish CD4+ and CD8+ T cells and NK cells from each other (Figure 2B). The B cells in our dataset were mostly plasma cells, expressing high levels of JCHAIN (Joining Chain of Multimeric IgA And IgM). IgM+ (IGHM) cells were enriched in the airway lumen and in the lung parenchyma, while IgG3+ (IGHG3) were enriched in airway biopsy samples and were virtually absent from the airway lumen. This suggests an isotype-driven micro-anatomical segregation of B cells in the airways (Extended Figure 5F).

### Molecular features of mucous cell metaplasia in asthma

To characterize the changes in the cellular landscape of airway wall in a chronic inflammatory condition, we also analysed bronchial biopsies from six volunteers with persistent, childhood-onset asthma (Figure 3A). Asthma is a complex disease^17^ and multiple cells such as epithelial^18,19^, endothelial^20^ and immune cells^21,22^ have been shown to be altered in asthma. The combined airway wall dataset reveals a cellular landscape dominated by epithelial (*EPCAM*-positive) cells, with minor contributions from endothelial, mesenchymal and immune cells (Extended Figure 6A and B).

**Figure 6.**
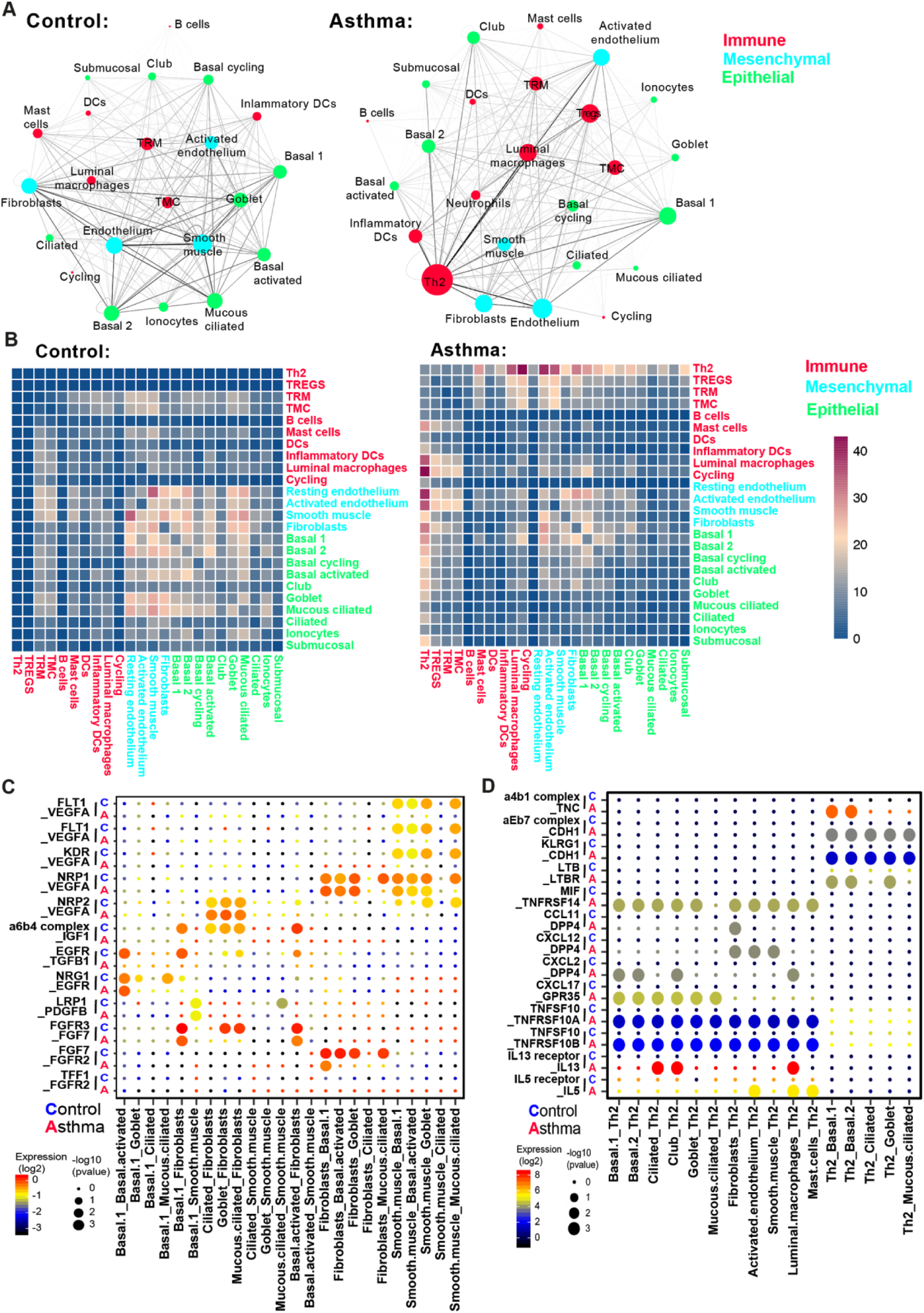
Asthma is characterized by unique cell-to-cell signalling networks. We quantified the predicted cell interactions in healthy and asthmatic airways between all the epithelial and non-epithelial cell clusters identified in figures 3 and 4, plus the lung airway enriched populations of CD4 T cells (Th2, Treg, TMC and TRM) **(A)** Networks depicting cell types as nodes and interactions as edges. Size of cell type proportional to the total number of interactions of each cell type and edge thickness proportional to the number of interactions between the connecting types. **(B)** Heatmap depicting the number of all possible interactions between the clusters analysed. Cell types grouped by broad lineage (epithelia, mesenchymal or immune). **(C)** Dot plot depicting selected epithelial-epithelial and epithelial-mesenchymal interactions enriched in healthy airways but absent in asthma. **(D)** Dot plot depicting selected epithelial-immune and mesenchymal-immune interactions highly enriched in asthma but absent in healthy airways.

High-resolution clustering of the EPCAM^+^ clusters identifies 10 sub clusters representing the 6 epithelial cell types observed in healthy airway wall (Figure 1C), as well as two additional basal cell states: a mucous ciliated cell state, and serous cells from the submucosal glands (Figure 3B). In addition to the two basal cell states observed in healthy airway wall (Figure 1C), the basal cell states in asthma include activated and cycling cell states (Figure 3B). Activated basal cells closely resemble Basal 1 cells in their transcriptional phenotype, but also express proinflammatory genes such as *POSTN* (Figure 3D). Cycling basal cells are characterized by expression of canonical marker genes of proliferating cells (MKI67 and TOP2A) (Figure 3D), and this is the only cluster of airway epithelial cells expressing the squamous cell marker *KRT13* (Extended Figure 6).

We observe mucous cell hyperplasia in asthma, with a strong increase in goblet cell numbers (Figure 3C), which are very rare in healthy airway wall biopsies (Figure 1E). Moreover, the goblet cell transcriptional phenotype is altered in asthma, with strongly increased expression of *MUC5AC* and *SPDEF*, as well as proinflammatory and remodelling genes including *NOS2*, CEACAM5 and *CST1* (Figure 3D). In addition, we identify a strong increase in mucous ciliated cells, a novel cell state that has remarkable transcriptional resemblance to ciliated cells, whilst co-expressing a number of mucous genes, including *MUC5AC, SERPINB2/3* and *CEACAM5* (Figure 3D, Extended Figure 6). Mucous ciliated cells lack expression of the transcription factor *SPDEF* (in contrast to club and goblet cells), while maintaining *FOXJ1* expression, underscoring their ciliated cell origin (Extended Figure 7).

To further dissect the inferred differentiation trajectories in healthy and asthmatic airway wall epithelial cells, we performed pseudotime analysis^23^. This reveals a trajectory starting with basal cell subsets, bifurcating into either a secretory lineage (mainly club cells) or a ciliated lineage in healthy airway wall (Figure 3E). In asthma, the secretory lineage is a mix of club and goblet cells, while the mucous ciliated cell state overlaps with the ciliated differentiation trajectory (Figure 3E,F).

Next, we further analysed the transcriptional profiles of the two mucous cell states we observe specifically in asthma: the mucous ciliated cells and the goblet cells. As both NOTCH and IL4/IL13 signalling have been shown to contribute to mucous cell differentiation^24^, we analysed expression of both NOTCH target genes^25,26^ and IL4/IL13 target genes^27^ in club, goblet, and ciliated cells as well as in the novel mucous ciliated cell state, in both asthma and healthy airway wall biopsies. Expression of IL4/IL13-induced genes^27^ is prominent in asthma, and highest in activated basal cells, goblet cells and mucous ciliated cells (Extended Figure 7). In club cells, expression of NOTCH target genes^25,26^ is not different between asthma and healthy-derived cells. In contrast, in goblet cells, the NOTCH target gene signature is retained only in cells from healthy airway wall, and is lost in asthma.

As in goblet cells, mucous ciliated cells also lack expression of Notch target genes in asthma (Extended Figure 7). Hence, we postulate that mucous ciliated cells represent a transition cell state in the ciliated lineage - induced by IL4/IL13 signalling - leading to a mucous cell phenotype which contributes to mucous cell metaplasia in asthma^24^. Similar to goblet cells, mucous ciliated cells express asthma genes such as *CST1*^*28*^ and *POSTN* (Figure 3D), indicating that these cells also contribute to airway inflammation and remodelling.

Integrating the asthma GWAS genes with our epithelial single cell transcriptomic data reveals a broad contribution of the airway epithelial cell types to asthma susceptibility (Figure 3G), with high expression of asthma GWAS genes in ciliated and mucous ciliated cells. This includes genes involved in cilia function (DYNC2H1 and KIF3A), cell adhesion (ELK3, CDHR3 and PTPRT) and IL5-induced mucus metaplasia (IL5RA)^29^, further suggesting a direct link between mucous ciliated cells and Th2 CD4 T cells.

### Remodelling of the stromal and Immune compartments in asthmatic airways

Asthma is associated with chronic inflammation and remodelling of the airway wall^30^. Analysis of the inflammatory and stromal cell populations in the bronchial biopsies by unsupervised clustering (Figure 4A) reveals the presence of B and T cells, neutrophils, macrophages, DCs, mast cells, fibroblasts, smooth muscle cells and endothelial cells (Figure 4B, Extended Figure 8). We did not detect any innate lymphoid cells, basophils or eosinophils as separate clusters (Extended Figure 8). Analysis of bulk transcriptome analysis of whole airway biopsies before and after tissue dissociation identified very low expression levels of eosinophil marker genes (CLC and IL5RA), indicating these cells are relatively rare in the samples we analysed (Extended Figure 9).

Mast cell numbers were increased in asthma (Figure 4C), while being virtually absent in the airways of healthy individuals (Figure 4C). Mast cells in asthmatic airways lack chymase 1 expression (*CMA1*) and express high levels of tryptase genes (*TPSB2, TPSAB1*) (Figure 4D). Prostaglandins and leukotrienes are known to be crucial to inflammatory cell signalling. Mast cells express high levels of PTGS2 and HPGDS (Figure 4D, Extended Figure 10). PTGS2 (cyclooxygenase-2), also known as inflammatory cyclooxygenase, converts the precursor arachidonic acid to prostaglandin endoperoxide H2 (PGH2). HPGDS (Hematopoietic Prostaglandin D Synthase) catalyses the conversion of PGH2 to prostaglandin D2 (PGD2). PGD2 activates CD4 Th2 cells^31^, ILC2^32^, basophils and neutrophils^31^ and plays a key role in asthma pathology. Expression of all PGD2 biosynthesis enzymes is a unique feature of mast cells (Extended Figure 10) and this suggests that intraepithelial mast cells are continually producing PGD2 in asthma patients. Thus, these cells are most likely intraepithelial mast cells, previously shown to accumulate in Th2-high asthmatic airway epithelium^33^, and reported to be increased^34^ with disease severity^21^.

We observed an increase in the number of B cells in the asthmatic airways (Figure 4C) and these cells have a plasma cell phenotype, with high JCHAIN expression (Figure 4D). The increase in B cell numbers was mostly of IgM+ cells (IGHM) (Figure 4E). IgM levels in asthma BALF samples have been reported to be either increased^34^ or unchanged ^35^, suggesting cohort dependent variability. IgM-producing B cells in the healthy airways were mostly present in the airway lumen (Extended Figure 5F). However, as we did not analyse brush samples from asthmatic patients, we cannot precisely pinpoint whether the increase in IgM+ B cells takes place in the intraepithelial region or in the lumen, as both regions are present in biopsy samples.

Asthma GWAS genes show highly cell-type restricted expression (Figure 4F). When excluding the widely expressed HLA genes from the analysis, fibroblasts and T cells express the highest number of asthma GWAS genes (Figure 4F), which are mostly upregulated in asthma (Figure 4F). GATA3 expression is restricted to T cells (Figure 4F), and increased in T cells from asthma patients (Figure 4F and G). We detected increased expression of CD4 (but not CD8a) in the T cell cluster, suggesting an increase in Th2 CD4 T cells (Figure 4G). Therefore, we next proceed to investigate the CD4 T cell compartment in depth.

### Pathogenic effector Th2 cells are enriched in asthmatic airways

In line with the increase in GATA3 and CD4 expression mentioned above, CD4 Th2 cells are known to be key drivers of asthma^17,36^. To assess the presence of Th2 effector cells in the airways of asthma patients (Figure 4G), we single cell sorted CD4 T cells followed by plate-based SmartSeq2 analysis for in depth transcriptional phenotyping of the T helper cell compartment (see Methods for details). We analysed cells from both peripheral blood and airway wall biopsies (Figure 5A) from a larger cohort of asthma patients and healthy controls (Figure 5B). Unbiased clustering reveals six major populations of CD4 T cells (Figure 5C and Extended Figure 11). At this coarse level of analysis, none of these six clusters was specifically enriched in asthma patients (Figure 5D).

Comparative analysis of CD4 T cells isolated from paired blood and lung samples allows us to differentiate between tissue-resident T cells and circulating T cells in an unbiased way, by subtracting the populations shared with blood from the populations specific to the biopsies (Figure 5E). Using this approach, we identified two subsets highly enriched in the lungs: the classical Tissue Resident Memory (TRM) CD4 T cells, and a novel subset we named Tissue Migratory CD4 T cell (TMC) (Figure 5E). Naive/central memory (CM), effector memory (EM), and EMRA cells, as well as a mixed Treg/Th2 cluster, are either enriched in blood or present in both blood and airways (Figure 5E).

To better understand the two distinct lung-restricted CD4 T cell subsets, we performed differential expression analysis between TRM and TMC cells (Figure 5F). Several transcription factors highly expressed in circulating cells are enriched in TMC cells, such as *LEF1, SATB1* and *KLF3*, while *ZEB2* is specific for TRM cells. TMC cells expressed the tissue egression markers *S1PR1*, CCR7 and *SELL* (CD62L) and lacked expression of the canonical TRM marker *ITGAE* (CD103) (Figure 5F). As low numbers of TMC cells were present in peripheral blood CD4 T cells (Figure 5E), these data suggest that these cells have the potential to transit between lung and blood, a hypothesis supported by pseudotime trajectory analysis of the CD4 T cell subsets (Extended Figure 12).

Protein expression of CD69 and CD103 (ITGAE) have both been used as hallmarks of lung resident CD4 T cells isolated from lung parenchyma^37,38^, but to the best of our knowledge, no similar analysis of lung airway epithelial CD4 T cells has been performed to date. Both TMC and TRM cells have high expression of CD69, but only TRM cells express ITGAE (Extended Figure 11), suggesting these subsets might be equivalent to the previously described cells^37,38^ However, TRM cells^37,38^ have been shown to lack S1PR1 and CCR7 protein expression, in contrast to TMC cells that express high mRNA levels of both markers. No direct whole transcriptome comparison of CD69+CD103+ *versus* CD69+CD103- TRMs has been performed yet^37,38^. Further studies are necessary to properly compare CD4 T cells from airway wall *versus* lung parenchyma, and investigate how TMC align with the previously reported TRM subsets at the transcriptome level.

TRM cells in airway wall expressed high levels of *CXCR6* and *ITGA1* and high levels of cytokines (*CCL4, CCL4L2, CCL5*) and effector molecules (*PRF1, GZMB, GZMA, GZMH*) (Figure 5F and Extended Figure 11), indicating they are also in a primed state capable of direct effector function, as recently shown for TRMs from lung parenchyma^37^.

CD4 effector T cells are classically divided into distinct functional subsets based on their cytokine profile^17^. We manually annotated clusters of Th1 (*IFNG*^+^), Th2 (*IL4*^+^, *IL5*^+^ or *IL13*^+^) and Th17 (*IL17A*^+^ or *IL17F*^+^) cells based on their cytokine expression profiles (Figure 5G, Extended Figure 13). Cytokine-producing cells were mostly retrieved from the biopsies, with only very few present in blood (Extended Figure 14).

In terms of absolute numbers, Th2 cells were very rare and found both in healthy and asthmatic patients, although numbers of Th2 cells were significantly increased in the airway wall in the asthma patients, with no detected difference in the relative proportions of the other subsets (Figure 5H). In addition to the signature cytokines and the transcription factor *GATA3* (Extended Figure 13), airway wall Th2 cells express *HGPDS*, identifying them as pathogenic effector Th2 (peTh2) cells, previously associated with eosinophilic inflammation of the gastrointestinal tract and skin^39^. Airway Th2 also express the transcription factor *PPARG*, and cytokine receptors *IL17RB* and *IL1RL1* (Figure 5I). *IL17RB* and *IL1RL1* have been reported as upregulated in pathogenic allergen-specific Th2 cells (coined Th2A cells), which are present in allergic disease^40^ as well as in chronic rhinosinusitis with nasal polyps^41^, suggesting airway wall Th2 cells share features with both Th2A and peTh2 cells.

### Asthma is characterized by specific signalling networks

Asthma is characterized by remodelling of the airways, which depends on complex interactions between structural and inflammatory cells^17^, both *via* direct physical interactions and *via* secreted proteins and small molecules. We used our recently developed receptor/ligand database and statistical inference framework CellPhoneDB^42^ (www.cellphonedb.org), to chart combinatorial cell-specific expression patterns of genes encoding receptor/ligand pairs. We aim to identify potential cell-cell interactions in the airway wall, and define their changes in asthma. Whilst most interactions are unchanged, some were specific to the diseased or healthy state (Full list in Extended 6).

In healthy controls, the cell-cell interaction landscape of the airway wall was dominated by lung structural cells (mainly mesenchymal and epithelial cell types) communicating with other lung structural cells and with tissue-resident T cells, both the classical TRM and the newly identified TMC subsets (Figure 6A,B, left panels). In the asthmatic airway wall, the number of predicted interactions between epithelial and mesenchymal cells was strongly reduced. Instead, the cell-cell communication landscape in asthmatic airway wall is dominated by Th2 cells that were found to have increased interactions with other immune cells, including antigen-presenting cells, and also with epithelial cells ((Figure 6A,B, right panels). The most striking increase in interactions is with mesenchymal cells, both fibroblasts and smooth muscle cells (Figure 6A and B, right panels).

Analysis of the predicted cell-cell interactions between structural cells in healthy airway wall revealed a wealth of growth factor-signalling pathways including the FGF, GFR, IGF, TGF, PDGF and VEGF pathways, most of which were lost in the asthmatic samples (Figure 6C). Detailed analysis of the individual interactions of Th2 cells with the inflammatory and lung structural cells unique to asthma revealed the potential of Th2 cells to have cognate interactions with the epithelial cells involving KLRG1 and CD103 binding to E-cadherin, and also integrin-mediated interactions with epithelial-expressed matrix proteins such as Tenascin-C. Epithelial expression of alarmins and cytokines, such as IL33, TSLP (Figure 3G) and TNFSF10/TRAIL (Figure 6D), all of which are asthma genes^43–45^, might then lead to activation of Th2 cells expressing the receptors.

In addition to validating these well-known interactions, which for IL33 and TSLP failed to reach significance in our unbiased cell-cell interaction analysis, we identify novel epithelial-Th2 cell interactions in asthma, including chemokines CXCL2 and CXCL17, and the cytokine MIF that are all expressed by epithelial cells, while Th2 cells express the respective receptors (Figure 6C). Interestingly, mesenchymal cells share some of these Th2 cell interactions, such as expression of TNFSF10/TRAIL and MIF. Predicted Th2 interactions unique to mesenchymal cells in asthmatic airway wall are CXCL12 and CCL11, expressed by fibroblasts and smooth muscle cells. Airway wall Th2 cells in asthma express the cytokines IL5 and IL13 (Figure 5I), the receptor complexes for which are expressed by immune cells and epithelial cells, respectively, in line with the expression of IL13-driven genes in mucous ciliated and goblet cells in asthma (Extended data Figure 7). In addition to these classical Th2 cytokines, Th2 cells express LTB for which basal epithelial cells express the receptor.

## Discussion

Our study profiles the cellular landscape of human lung tissue at the single-cell level, including both upper and lower airways and parenchymal lung tissue in healthy adults. We identify at least 21 main cell types in the normal human lung, that can be further subdivided into more fine-grained cell states. There is clear Mendelian disease relevance for many cells, including the previously-reported ionocytes (for bronchiectasis and cystic fibrosis), and type-2 alveolar cells (for alpha-1-antitrypsin deficiency). We chart differences in frequencies and molecular state of airway epithelial cells between upper and lower airways. To our knowledge, we provide for the first time a detailed molecular description of tissue-resident CD4 T cells in the human lower airway wall, and identify two separate subsets, one of which was hitherto unknown.

In our studies, we deployed two different droplet-based single-cell RNA sequencing platforms and one plate-based method, with experiments performed in three geographically distinct research centers. Overall, the datasets are remarkably consistent, which yields a quantitative dataset of the cellular composition of human upper and lower airways and lung parenchyma.

In addition to analysing healthy reference samples, we characterize the changes in the cell types and cell states in airway wall in asthma. This reveals mucous ciliated cells as a novel cell state that contributes to mucous cell metaplasia. Both mucous ciliated and goblet cells are characterized by expression of an IL4/IL13-driven gene signature, indicating a dominant role for type-2 cytokines in maintaining the epithelial changes in chronic airway inflammation in asthma. The mucin gene expression induced in FOXJ1+ cells with an unabated ciliated cell transcriptional profile strongly indicates that the mucous state is superimposed on the ciliated cell phenotype, independent of goblet cell differentiation from club cells. Our data seem to indicate that these two processes can occur in parallel, with mucous metaplasia of ciliated cells and goblet cell hyperplasia both contributing to the increase in mucin-producing cells in asthma. Whether the mucous ciliated cells go on to lose their FOXJ1 expression and the ciliated transcriptional profile and transdifferentiate into *bona fide* SPDEF-positive goblet cells remains to be firmly established.

The epithelial cell changes in airway wall in asthma are surprisingly different to those recently described in patients with in chronic rhinosinusitis with polyps^27^. In this chronic inflammatory disease of the upper airways caused by exaggerated type-2 immunity, an IL4/IL13-driven gene transcription profile was mainly observed in basal epithelial cells, which were found to be arrested in differentiation and highly increased in numbers^27^. Using the same IL4/IL13-driven gene module, we find some expression thereof in basal cells, but in asthma this does not result in significant changes in basal cell numbers. Instead, we observe an increased number of goblet cells, as well as of mucous ciliated cells, both of which show evidence of marked expression of the IL4/IL13-driven gene signature (extended data Figure 7). Hence, while there is some overlap in the cellular mechanisms underlying rhinosinusitis with polyps and asthma, the resultant changes in cellular states and their frequencies in airway wall differ considerably between the epithelia of the upper *versus* the lower airways. In contrast, the changes in the eicosanoid pathway observed in chronic type-2 inflammation of the upper^27^ and lower (extended data Figure 10) airways are very similar, likely reflecting a common cellular mechanism between Th2 inflammation in these two anatomical locations.

Conflicting data have been reported on dependence of IL13-induced goblet cell metaplasia on NOTCH signalling in *ex vivo* cultured PBECs^25,46^. Our data on freshly isolated bronchial epithelial cells show that expression of NOTCH2 and Notch target genes is present in goblet cells from healthy controls only, and absent from ciliated, mucous ciliated and goblet cells in asthmatic airway wall. This indicates that mucous cell metaplasia in mild asthmatics is likely to be driven by type-2 cytokines in a NOTCH-independent fashion. One likely source of the type-2 cytokines driving this goblet cell metaplasia are the cytokine-expressing effector Th2 cells that are increased in asthmatic airway wall. To our knowledge, our study is the first to conclusively show the presence of the recently identified^39,40,41^ pathogenic effector Th2 cells in the airway wall in asthma, as evidenced by the combined expression of *IL5, IL13, HPGDS, PPARG, IL17RB, IL1RL1* and *IL1RAP*. Additional cellular sources for the type-2 cytokines, including innate lymphoid cells, cannot be ruled out based on our data, as these cells were present in too low numbers to be analysed in our dataset, and will need to be purified from the biopsy cell suspensions for further characterization in future studies.

Finally, our detailed and unbiased analyses of cell-cell communication in healthy and asthmatic airway wall reveals novel interactions of the airway wall resident cells in health and disease. Comprehensive analysis of the cell-cell interactions underpinning the changes of the airway wall cellular landscape in asthma identifies a shift away from interactions between structural cells in healthy airway wall, towards an intercellular network dominated by the interactions of Th2 cells with structural and inflammatory cells in asthmatic airway wall. The richness of growth factor signalling between epithelial cells and mesenchymal cells observed in healthy airway wall is largely lost in asthma, which seems at odds with a reactivation of the epithelial-mesenchymal trophic unit in asthma^47^. Instead, our data supports a shift in cellular phenotypes in airway wall due to the local production of Th2 cytokines such as IL13 in chronic disease in our patient cohort with childhood-onset asthma. This global view of the airway wall cellular landscape in health and in asthma opens up new perspectives on lung biology and molecular mechanisms of asthma.

## Acknowledgments

We thank Jana Elias (scientific illustrator) for the design of cartoons, the Sanger Single Cell Genomics Core Facility for support with the SmartSeq2 protocol, Dr. Emma Rawlins for feedback and critical reading of the manuscript, as well as all the members of the Teichmann lab for scientific input. We are grateful to the Cambridge Biorepository for Translational Medicine for the provision of tissue from deceased organ donors.

## Authors contribution

*<u>Designed the project:</u>* Teichmann, S.A, Nawijn, M.C., van den Berge, M., Affleck, K., van Oosterhout, A.J., Schiller, H.B. *<u>Wrote the paper:</u>* Vieira Braga, F.A., Nawijn, M.C., Teichmann, S.A. *<u>Generated data:</u>* Vieira Braga, F.A., Carpaij, O.A., Brouwer, S., Hesse, L., Jiang, J., Fasouli, E.S., Strzelecka, P., Mahbubani, K.T.A., Angelidis, I., Strunz, M. *<u>Analysed</u> <u>data:</u>* Vieira Braga, F.A., Kar, G., Berg, M. Simon, L., Gomes, T., Jiang, J., Efremova, M., Palit, S., Polanski, K., Firth, H.V., Theis, F.J. *<u>Interpreted data:</u>* Vieira Braga, F.A., Kar, G., Carpaij, O.A., Simon, L., Gomes, T., Jiang, J., Vento-Tormo, R., Affleck, K., Palit, S., Cvejic, A., Saeb-Parsy, K., Timens, W., Koppelman, G.H., van Oosterhout, A.J., Schiller, H.B., van den Berge, M., Theis, F.J., van den Berge, M., Meyer, K.B. All authors read the manuscript, offered feedback and approved it prior to submission.

## Funding

This work was funded by Open Targets, an open innovation public-private partnership (http://www.opentargets.org), a GlaxoSmithKline collaborative agreement with University Medical Center Groningen, Wellcome (WT206194), EMBO and HFSP Long Term fellowships to R. Vento-Tormo, the Marie Curie ENLIGHT-TEN training network for Tomas Gomes, the Lung Foundation Netherlands (projects no 5.1.14.020 and 4.1.18.226), and Health-Holland, Top Sector Life Sciences & Health. LMS acknowledges funding from the European Union’s Horizon 2020 research and innovation programme under the Marie Sklodowska-Curie grant agreement No 753039.

## Competing interests

Affleck, K and van Oosterhout, A.J are employees of GSK.

## Data availability

Interactive exploration tool: www.lungcellatlas.org

## Methods

### Patient recruitment and ethical approval

#### Bronchoscopy biopsy (10x and smartseq2 analysis)

cohort inclusion criteria for all subjects were: age between 40 – 65 years and history of smoking <10 pack years. For the asthmatics, inclusion criteria were: age of onset of asthmatic symptoms ≤12 years, documented history of asthma, use of inhaled corticosteroids with(out) β2-agonists due to respiratory symptoms and a positive provocation test (i.e. PC20 methacholine ≤8mg/ml with 2-minute protocol). For the non-asthmatic controls, the following criteria were essential for inclusion: absent history of asthma, no use of asthma-related medication, a negative provocation test (i.e. PC20 methacholine >8 mg/ml and adenosine 5’-monophosphate >320 mg/ml with 2-minute protocol), no pulmonary obstruction (i.e. FEV1/FVC ≥70%) and absence of lung function impairment (i.e. FEV1 ≥80% predicted).

Asthmatics stopped inhaled corticosteroid use 6 weeks prior to all tests. All subjects were clinically characterised with pulmonary function and provocation tests, blood samples were drawn, and finally subjects underwent a bronchoscopy under sedation. If a subject developed upper respiratory symptoms, bronchoscopy was postponed for ≥6 weeks.

Fibreoptic bronchoscopy was performed using a standardised protocol during conscious sedation [1]. Six macroscopically adequate endobronchial biopsies were collected for this study, located between the 3^rd^ and 6^th^ generation of the right lower and middle lobe. Extracted biopsies were processed directly thereafter, with a maximum of one hour delay. The medical ethics committee of the Groningen University Medical Center Groningen approved the study, and all subjects gave their written informed consent. Detailed patient information below:

| Donor | Gender | Age | Classification | Packyears | BMI | FEV1% pred | FEV1/FVC ratio | PC20 methacholine threshold (mg/ml) | 10x | Smartseq2 |
| --- | --- | --- | --- | --- | --- | --- | --- | --- | --- | --- |
| ARMS004 | F | 57 | Ctrl | 0 | 21 | 99 | 82 | Not reached | no | yes |
| ARMS005 | F | 44 | Ctrl | 4 | 24 | 111 | 82 | Not reached | no | yes |
| ARMS009 | M | 49 | Asthma | 0 | 22 | 87 | 69 | 0.211 | no | yes |
| ARMS014 | F | 65 | Ctrl | 0 | 18 | 109 | 78 | Not reached | no | yes |
| ARMS015 | M | 49 | Asthma | 0 | 24 | 105 | 67 | 1.155 | no | yes |
| ARMS018 | M | 54 | Ctrl | 3 | 25 | 107 | 80 | Not reached | no | yes |
| ARMS019 | F | 56 | Asthma | 0 | 45 | 44 | 55 | FEV1 too low | no | yes |
| ARMS022 | M | 48 | Ctrl | 1 | 30 | 131 | 84 | Not reached | no | yes |
| ARMS024 | F | 51 | Asthma | 0 | 33 | 96 | 74 | 0.88 | yes | yes |
| ARMS026 | M | 64 | Ctrl | 5 | 32 | 103 | 81 | Not reached | yes | yes |
| ARMS032 | M | 56 | Ctrl | 0 | 22 | 120 | 75 | Not reached | yes | yes |
| ARMS033 | M | 51 | Asthma | 0 | 23 | 98 | 69 | 2.63 | yes | yes |
| ARMS035 | M | 61 | Asthma | 0 | 20 | 89 | 59 | 6.71 | yes | yes |
| ARMS038 | M | 59 | Ctrl | 0 | 23 | 148 | 74 | Not reached | yes | yes |
| ARMS040 | M | 51 | Ctrl | 0 | 23 | 119 | 73 | Not reached | yes | yes |
| ARMS043 | F | 50 | Ctrl | 1 | 23 | 110 | 71 | Not reached | yes | no |
| ARMS048 | M | 53 | PersA | 0 | 29 | 77 | 54 | 0.205 | yes | no |
| ARMS050 | M | 64 | PersA | 0 | 21 | 85 | 66 | 0.692 | yes | no |
| ARMS051 | M | 64 | PersA | 0 | 25 | 59 | 54 | 0.631 | yes | no |
| ARMS054 | M | 60 | Ctrl | 0 | 29 | 116 | 81 | Not reached | yes | no |

#### Lung resection (Dropseq analysis)

Fresh resected human lung tissue (parenchymal lung and distal airway specimens) was obtained via the CPC BioArchive at the Comprehensive Pneumology Center Munich (CPC-M, Munich, Germany). In total, we analysed parenchymal tissue of uninvolved areas of tumour resection material from four patients. All participants gave written informed consent and the study was approved by the local ethics committee of the Ludwig-Maximilians University of Munich, Germany.

For transport from the surgeon to the laboratory, lung tissue samples were stored in ice-cold DMEM-F12 media and packed in thermo stable boxes. Tissue was processed with a maximum delay of 2 hours after surgery. Upon delivery to the lab, tissue samples were assessed visually for qualification for the study.

| ID | Sample | Gender | Age | Smoker | Segment | COPD |
| --- | --- | --- | --- | --- | --- | --- |
| muc3843 | ASK428 | m | 62 | yes,<br>>100py | lower lobe, left | COPD Gold II |
| muc4658 | ASK440 | m | 58 | yes, 45py | upper lobe, left | no |
| muc5103 | ASK452 | m | 58 | no | main bronchus | no |
| muc5104 | ASK454 | m | 69 | yes, >60py | upper lobe, left | no |

#### Lung transplant tissue (10x analysis)

Human lung tissue was obtained from deceased organ donors from whom organs were being retrieved for transplantation. Informed consent for the use of tissue was obtained from the donors’ families (REC reference: 15/EE/0152 NRES Committee East of England-Cambridge South).

Fresh tissue from the peripheral parenchyma of the left lower lobe or lower right lobe of the lung was excised within 60 minutes of circulatory arrest and preserved in University of Wisconsin (UW) organ preservation solution (Belzer UW® Cold Storage Solution, Bridge to Life, USA) until processing.

Donor information:

*Donor 284C*

Gender: Male

Age band: 55-60

BMI: 25.83

Cause of Death: hypoxic brain damage

Smoking history: smoked 20/day for 25 years

Stopped: 08/2000

Respiratory related information: Chest X-ray normal on admission. No pleural effusion or pneumothorax. Not diagnosed with asthma, but inhalers for possible seasonal wheeze. Family report only using inhaler approximately 5 times a year. No recent peak flow on record last one in 2008 when it was 460, predicted is 611. Time from death to cell lysis: 12h

*Donor 290B*

Gender: Female

Age band: 60-65

BMI: 27.77

Cause of Death: hypoxic brain damage

Smoking history: smoked 15/day for 7 years

Stopped: no details

Respiratory related information: Respiratory tests all normal on admission; maintaining own airway. GP notes report Acute bronchitis in 1994. Time from death to cell lysis: 2h 27min

*Donor: 292B*

Gender: Male

Age band: 55-60

BMI: 27.44

Cause of Death: Intracranial haemorrhage

Smoking history: smoked 20/day for 46 years

Stopped: no details

Respiratory related information: Chest X-ray normal on admission, lungs appear clear. Bronchoscopy results show global inflamed mucosa. No other history of respiratory issues. Time from death to cell lysis: 18h 50min

*Donor: 296C*

Gender: Female

Age band: 30-35

BMI: 20.9

Cause of Death: Intracranial haemorrhage

Smoking history: smoked 20/day for 19 years

Stopped: no details

Respiratory related information: Chest X-ray shows collapsed left lobe on admission due to consolidation. Right lobe looks normal. No history or record of respiratory issues. Time from death to cell lysis: 15h 30min

*Donor: 298C*

Gender: Male

Age band: 50-55

BMI: 24

Cause of Death: Intracranial haemorrhage

Smoking history: not available

Stopped: no details

Respiratory related information: no details Time from death to cell lysis: 15h 30min

*Donor: 302C*

Gender: Male

Age band: 40-45

BMI: 34.33

Cause of Death: Known or suspected suicide

Smoking history: smoked 20/day for 25 years

Stopped: no details

Respiratory related information: Chest X-ray shows reduced volume in right lung due to collapsed right lower lobe on admission. No history or record of respiratory issues. Time from death to cell lysis: 13h 30min

#### Archived formalin-fixed paraffin-embedded (FFPE) lung blocks

Left-over frozen peripheral lung tissues from 6 current smokers and 4 non-smokers who underwent lung resection surgery. These subjects did not have a history of lung disease, apart from lung cancer for which the patients underwent surgery. Lung tissue samples were taken as distant from the tumor as possible. Thus, any possible effect of the tumor on the lung tissue was minimized. All samples were obtained according to national and local ethical guidelines and the research code of the University Medical Center Groningen.

| *Bio sample code* | *Age* | *Gender* | *Smoking status* | *Diagnosis* |
| --- | --- | --- | --- | --- |
| GRNG-LNG_MRK085 | 45 | female | current smoker | LCC |
| GRNG-LNG_MRK125 | 54 | female | current smoker | AC |
| GRNG-LNG_MRK177 | 52 | female | current smoker | AC |
| GRNG-LNG_MRK188 | 70 | male | current smoker | SCC |
| GRNG-LNG_MRK218 | 61 | male | current smoker | Metastasis |
| GRNG-LNG_MRK245 | 65 | female | current smoker | LCC |
| GRNG-LNG_0758 | 81 | female | non-smoker | AC |
| GRNG-LNG_0764 | 49 | male | non-smoker | carcinoid |
| GRNG-LNG_0802 | 50 | male | non-smoker | metastasis |
| GRNG-LNG_0830 | 45 | male | non-smoker | IMT |

LCC = large cell carcinoma, AC = adenocarcinoma, SCC = squamous cell carcinoma and IMT = inflammatory myofibroblast tumor

### Blood processing

Lithium Heparin-anticoagulated whole blood (500µl) was lysed using an ammonium chloride-potassium solution (155mM ammonium chloride (NH4Cl), 10mM potassium bicarbonate (KHCO3), 0,1mM EDTA). Cells were centrifuged for 5 min at 4°C, 550g after which the cell pellet was washed twice with PBS containing 1% BSA, followed by staining for cell surface markers.

### Lung tissue processing

*Bronchoscopy biopsy:* A single cell solution was obtained by chopping the biopsies finely using a single edge razor blade. The chopped tissue was then put in a mixture of 1mg/ml collagenase D and 0.1mg/ml DNase I (Roche) in HBSS (Lonza). This was then placed at 37°C for 1hr with gentle agitation. The single cell suspension was forced through a 70µm nylon cell strainer (Falcon). The suspension was centrifuged at 550g, 4°C for 5 min and washed once with a PBS containing 1% BSA (Sigma Aldrich). The single cell suspensions used for 10X Genomics scRNAseq analysis were cleared of red blood cells by using a Red blood cell lysis buffer (eBioscience) followed by staining for cell surface markers.

*Lung tissue resection:* For each sample, 1-1.5 g of tissue was homogenized by mincing with scissors into smaller pieces (∼0.5 mm2/piece). Prior to tissue digestion, lung homogenates were cleared from excessive blood by addition of 35 ml of ice-cold PBS, followed by gentle shaking and tissue collection using a 40μm strainer. The bloody filtrate was discarded. The tissue was transferred into 8 ml of enzyme mix consisting of dispase (50 caseinolytic U/ml), collagenase (2 mg/ml), elastase (1 mg/ml), and DNase (30 µg/ml) for mild enzymatic digestion for 1 hour at 37°C while shaking. Enzyme activity was inhibited by adding 5 ml of PBS supplemented with 10% FCS. Dissociated cells in suspension were passed through a 70μm strainer and centrifuged at 300g for 5 minutes at 4°C. The cell pellet was resuspended in 3 ml of red blood cell lysis buffer and incubated at room temperature for 2 minutes to lyse remaining red blood cells. After incubation, 10 ml of PBS supplemented with 10% FCS was added to the suspension and the mix was centrifuged at 300g for 5 minutes at 4°C. The cells were taken up in 1 ml of PBS supplemented with 10% FCS, counted using a Neubauer chamber and critically assessed for single cell separation. Dead cells were counted to calculate the overall cell viability, which needed to be above 85% to continue with Drop-Seq. 250,000 cells were aliquoted in 2.5 ml of PBS supplemented with 0.04% of bovine serum albumin and loaded for Drop-Seq at a final concentration of 100 cells/μl.

*Rejected lung transplant:* for each sample, 1-2g of tissue was divided in 5 smaller pieces then transferred to 5ml eppendorfs containing 1.5ml 0.5mg/ml collagenase D and 0.1mg/ml DNase I (Sigma) in RPMI. Samples were then finely minced using scissors. Minced tissue was then transferred to a petry dish and extra digestion medium added to completely cover the tissue. Samples were incubated 30min at 37°C. Cells were then passed up and down through a 16-gauge needle 10 times. Samples were incubated for an additional 15min at 37°C. Cells were filtered a 70um filter, then spun down for 6min 1400RPM. 1 ml of red blood cell lysis (eBioscience) was added to the pellet during 5min. Cells were resuspended in RPMI + 10%FCS and counted. Dead cells were removed using the Dead Cell Removal Kit (Miltenyi Biotec). In brief, cells were incubated with anti-Annexin V beads for 15min. The cell suspension was then passed through a magnetic column and dead Annexin V+ cells remained in the column, while live cells were collected. Viability was then estimated *via* trypan blue. More than 99% of cells were viable.

### Flow cytometry

Blood leukocytes were stained with CD4 APC-Cy7, CD3 PerCP Cy5.5 and CD8 APC (eBioscience) for 30min at 4°C and washed twice with PBS containing 1% BSA. Propidium iodide (PI) was added 5min before sorting.

Airway wall biopsy single cell suspensions were stained for 30min at 4°C with CD3 PerCP Cy5.5, CD45 BB515, CD4 APC Cy7 (BD) and CD8 PE and washed twice with PBS containing 1% BSA. Propidium iodide (IQ products) was added 5min before sorting.

### Cell Sorting

Lymphocytes were selected in the FCS/SSC plot. These were then selected on single, live cells for blood or single, live, CD45+ for lung. The sorted cells were positive for CD3 & CD4 as shown in figure 5A. All cells were sorted in a MoFlo Astrios (Beckman Coulter) using Summit Software (Beckman Coulter).

### Immunohistochemical staining

Human lung tissue containing large airways were collected from archival formalin-fixed paraffin-embedded (FFPE) blocks (n=10, 6 smokers and 4 non-smokers). Serial sections (∼4 µm) were cut for immunohistochemistry (IHC) and immunofluorescent (IF) staining.

Serial sections from FFPE lung tissue were stained for using standard protocols, with antibodies specified in the figures. Briefly, serial sections were deparaffinized in xylene, rehydrated and immersed in 10 mM sodium citrate buffer (pH 6.0). Antigen retrieval was performed by boiling the sections in a pressure cooker at 120°C for 20 min.

IHC and IF staining was performed as described previously^49,50^. For the IHC staining cells were stained with a primary antibody (see below for Ab details) and visualized with diaminobenzidine (DAB, Sigma) solution. For the IF staining, cells were stained with primary antibody. Secondary antibodies conjugated to fluorophores (donkey anti rabbit-488, donkey anti mouse-555) were used at a dilution of 1:100. DAPI, dissolved in Dako Fluorescence Mounting Medium (Dako S3023) at a dilution of 1:1000, was used as a nuclear stain.

### Antibody list

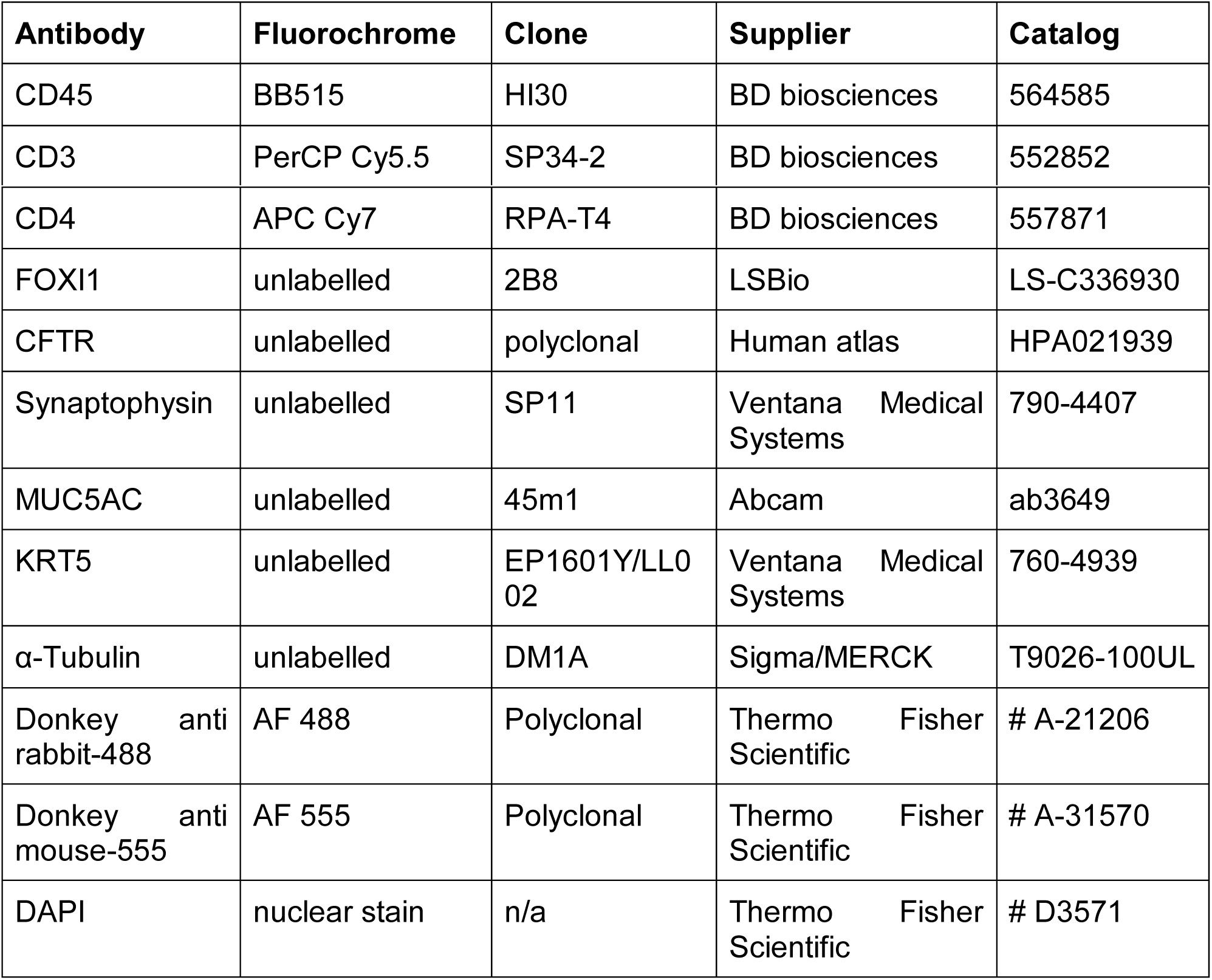

### Chromium 10x Genomics library and sequencing

#### Airway biopsy

Single cell suspensions were manually counted using a haemocytometer and concentration adjusted to a minimum of 300 cells/ul. Cells were loaded according to standard protocol of the Chromium single cell 3’ kit in order to capture between 2000-5000 cells/chip position. All the following steps were performed according to the standard protocol. Initially, we used one lane of an Illumina Hiseq 4000 per 10x Genomics chip position. Additional sequencing was performed in order to obtain coverage of at least mean coverage of 100.000 reads/cell.

#### Lung transplant

Single cell suspensions were manually counted using a haemocytometer and concentration adjusted to 1000 cells/ul. Cells were loaded according to standard protocol of the Chromium single cell 3’ kit in order to capture between 2000-5000 cells/chip position. All the following steps were performed according to the standard manufacturer protocol. Initially, we used one lane of an Illumina Hiseq 4000 per 10x Genomics chip position. Additional sequencing was performed in order to obtain coverage of at least mean coverage of 100.000 reads/cell.

### SmartSeq 2 library preparation and sequencing

Library preparation was performed with minor modifications from the published SmartSeq2 protocol^51^. In short, single cells were flow sorted onto individual wells of 96 or 384 wells containing 4ul (96 wells) or 1ul (384 wells) of lysis buffer (0.3% triton plus DNTPs and OligoDT). After sorting, plates were frozen and stored at −80 until further processing. RT, PCR (25 cycles) and nextera library preparation performed as described in ^51^.

### Dropseq library preparation and sequencing

Drop-seq experiments were performed largely as described previously^7^ with few adaptations during the single cell library preparation. Briefly, using a microfluidic polydimethylsiloxane (PDMS) device (Nanoshift), single cells (100/µl) from the lung cell suspension were co-encapsulated in droplets with barcoded beads (120/µl, purchased from ChemGenes Corporation, Wilmington, MA) at rates of 4000 µl/hr. Droplet emulsions were collected for 15 min/each prior to droplet breakage by perfluorooctanol (Sigma-Aldrich). After breakage, beads were harvested and the hybridized mRNA transcripts reverse transcribed (Maxima RT, Thermo Fisher). Unused primers were removed by the addition of exonuclease I (New England Biolabs), following which beads were washed, counted, and aliquoted for pre-amplification (2000 beads/reaction, equals ∼100 cells/reaction) with 12 PCR cycles (primers, chemistry, and cycle conditions identical to those previously described. PCR products were pooled and purified twice by 0.6x clean-up beads (CleanNA). Prior to tagmentation, cDNA samples were loaded on a DNA High Sensitivity Chip on the 2100 Bioanalyzer (Agilent) to ensure transcript integrity, purity, and amount. For each sample, 1 ng of pre-amplified cDNA from an estimated 1000 cells was tagmented by Nextera XT (Illumina) with a custom P5 primer (Integrated DNA Technologies). Single cell libraries were sequenced in a 100 bp paired-end run on the Illumina HiSeq4000 using 0.2 nM denatured sample and 5% PhiX spike-in. For priming of read 1, 0.5 µM Read1CustSeqB (primer sequence: GCCTGTCCGCGGAAGCAGTGGTATCAACGCAGAGTAC) was used.

### Bulk Transcriptome

Biopsies were fresh frozen in liquid nitrogen and stored in −80. RNA was extracted after a few weeks using a combination of Trizol & the RNeasy MinElute Clean Up kit from Qiagen. RNA was prepared from sequencing using the TruSeq RNA Library Prep Kit v2. Samples were then sequenced inn a Hiseq 4000.

### Single-cell RNA sequencing data alignment

For SmartSeq2 raw sequencing data, paired-end reads were mapped to the Human genome (GRCh38) using GSNAP with default parameters^52^. Then, uniquely mapped reads were counted using htseq-count (http://www-huber.embl.de/users/anders/HTSeq/). Low-quality cells were filtered out using the outlier detection algorithm in R Scater package based on a cut-off of 2*MAD (median-absolute-deviation).

10X Genomics raw sequencing data was processed using CellRanger software version 2.0.2 and the 10X human genome GRCh38 1.2.0 release as the reference.

The Dropseq core computational pipeline was used for processing next generation sequencing reads of the Dropseq scRNA-seq data, as previously described^7^. Briefly, STAR (version 2.5.2a) was used for mapping^53^. Reads were aligned to the human reference genome hg19 (provided by Dropseq group, GSE63269).

### Bulk transcriptome computational analysis

The bulk samples were aligned using STAR 2.5.1b, using the STAR index from the GRCh38 reference that was used when mapping 10X data, and quantified using HTSeq. The data was then processed using the Seurat-inspired workflow within Scanpy, adding a number of “pseudo-bulks” obtained by taking 10X data from donors matching the bulk samples and summing expression across all cells.

### Data QC

#### General strategy for 10x datasets

Optimal tissue dissociation conditions are cell-type dependent, resulting in a certain degree of cell lysis when working with a mixed tissue sample. This results in substantial background levels of ambient RNA in the single-cell suspension that vary with cell type composition, so we applied SoupX for background correction (see below). We analysed each donor sample separately and excluded cells with a number of genes higher than the median+2SDs for that donor. We further excluded cell with high number of UMIs and high percentage of mitochondrial reads (see below).

In parallel, we used scrublet (see below) to infer the number of the doublets in the dataset before applying the filters previously described and excluded any remaining cells predicted to be doublets that were still present in the dataset. We normalised and scaled our data (see below), performed clustering (see below) and identified and subset the data into epithelial and non-epithelial cell groups (as shown in supplementary figures 1 and 6). After separation between epithelial and non-epithelial, we clustered the cells and performed curated doublet removal (see below) based on known lineage restricted markers.

#### General strategy for Dropseq data

We normalised and scaled the data, then performed filtering based on the number of genes and percentage of mitochondrial reads.

#### General strategy smartseq2 data

We normalised and scaled the data, then performed filtering based on the number of genes and percentage of mitochondrial reads. In order to avoid potential batch effects from the lung digestion protocol, we corrected the gene expression of the CD4 SmartSeq2 dataset using a small subset of genes the expression of which has been recently shown to be highly responsive to enzymatic digestion^54^: FOS, ZFP36, JUN, FOSB, HSPA1A, JUNB, EGR1, UBC.

### Ambient RNA correction (SoupX)

Different batches can be affected by different levels of ambient RNA. To take this into account, we used the recently developed SoupX method^55^. Briefly, ambient RNA expression is estimated from the empty droplet pool (10 UMI or less). Expression of these genes in each cell is then calculated and compared to their proportion in the ambient RNA profile. Transcripts with a bimodal profile (i.e. that characterize specific groups of cells but are also highly abundant in empty droplets) are then grouped based on their function. The contamination fraction derived from the expression of these genes is then used to calculate the fraction of each droplet’s expression corresponding to the actual cell. Finally, this fraction and the ambient profiles are subtracted from the real expression values.

### UMI and number of genes filtering

#### 10x data (After SoupX correction)

nUMI: minimum 1000/ maximum 60000.

percent.mito, minimum 0 / maximum= 3%

#### SmartSeq2 data

nGene: minimum 1000 / maximum 4000.

percent.mito, minimum 0 / maximum= 15%

#### Dropseq data

nGene: minimum 200/ maximum 4000.

percent.mito, minimum 0 / maximum= 20%

### Scrublet

We used Scrublet (Wolock et al, BioRxiv, https://doi.org/10.1101/357368) for unbiased computational doublet inference. Doublets were identified in each 10X sample individually using scrublet, setting the expected doublet rate to 0.03 and keeping all other parameters at their default values. Cells were excluded when they had a score higher than 0.1 for upper and lower airway samples or higher than 0.05 for parenchyma samples.

### Normalisation and scaling

Downstream analyses including, normalisation, scaling, clustering of cells and identifying cluster marker genes were performed using the R software package Seurat^48^ version 2.1 (https://github.com/satijalab/seurat).

Samples were log normalised and scaled for the number of genes, number of UMIs and percentage of mitochondrial reads. The epithelial biopsy dataset comparing healthy and asthma was also scaled for XIST expression, as we observed some gender specific clusters of cells that shared lineage markers with the other observed clusters.

### Curated doublet removal

We combined literature knowledge about cell lineages with over clustering to identify clusters enriched in potential doublets. The strategy for each dataset is shown below:

#### *Lung atlas epithelial dataset (Figure 1* and associated extended data figures)

We removed cells with expression level higher than 0.5 for any of the following markers: PTPRC (immune),FCER1G (immune), PDGFRA (fibroblast) or PECAM1 (endothelial).

#### *Lung atlas non-epithelial dataset (Figure 2* and associated extended data figures)

We removed cells with expression level higher than 0.5 for any of the following markers: EPCAM (epithelial),KRT5 (basal), “FOXJ1”(ciliated) or MUC5AC (secretory). We then performed first clustering round (7 PCs, resolution 2) and excluded clusters that expressed combinations of the following lineage specific markers: MARCO(macrophage), CCL21 (lymphatic endothelial), TPSB2 (mast cell) or CD3D(T cell). We performed a second clustering round and exclude a cluster formed by cells from one donor that had low expression TPSB2, while lacking markers for all other immune lineages.

#### *Asthma biopsy epithelial cells* (Figure 3 and associated extended data figures)

due to the smaller number of cells, we only performed cluster-based doublet exclusion, without cell filtering. We performed one round of clustering and removed one clusters with high expression of PECAM1 (endothelial marker).

#### *Asthma biopsy non-epithelial cells* (Figure 4 and associated extended data figures)

we performed three rounds of clustering where we excluded clusters with high levels of EPCAM or KRT5 expressed in much higher levels than immune lineage markers.

### Dimensionality reduction

We performed PCA dimensionality reduction with the highly variable genes as input. We then used the PCs to calculate t-Distributed Stochastic Neighbour Embedding (**t-SNE**) for each dataset, using a perplexity value of 50.

### Data clustering

We used the function “FindClusters” from Seurat. In brief, this method uses a shared nearest neighbour (SNN) modularity optimization-based clustering algorithm to identify clusters of cells based on their PCs. Before constructing the SNN graph, this function calculates k-nearest neighbours (we used k=30) and then it constructs the SNN graph. The number of PCs used for each clustering round was dataset dependent and they were estimated by the elbow of a PCA scree plot, in combination to manual exploration of the top genes from each PC.

### DE analysis

We used a Wilcoxon rank sum test to identify differentially expressed genes in all the comparisons here discussed.

### MatchScore

We used MatchSCore^8^ to quantify the overlap of cell type marker signatures between experiments, which is based on the Jaccard index. Only marker genes with adjusted p-value < 0.1 and average log fold change > 1 were considered.

### CellPhoneDB

We developed a manually curated repository of ligands, receptors and their interactions called CellPhoneDB (www.cellphonedb.org; Vento-Tormo, Efremova et al., *Nature*, 2018), integrated with a statistical framework for predicting cell-cell communication networks from single cell transcriptome data. Briefly, the method infers potential receptor-ligand interactions based on expression of a receptor by one cell type and a ligand by another cell type. Only receptors and ligands expressed in more than 30% of the cells in the specific cluster were considered. In order to identify the most relevant interactions between cell types, the method prioritizes ligand-receptor interactions that have cell type-specific expression. To this end, pairwise cluster-cluster interaction analysis are performed by randomly permuting the cluster labels of each cell 1000 times. For each permutation, the total mean of the average receptor expression level of a cluster and the average ligand expression level of the interacting cluster is calculated, and a null distribution is derived for each receptor-ligand pair in each cluster-cluster interaction. An empirical *p*-value is calculated from the proportion of the means which are “as or more extreme” than the actual mean. For the multi-subunit heteromeric complexes, the member of the complex with the minimum average expression is used for calculating the mean.

Network visualization was done using Cytoscape (version 3.5.1). All the interaction pairs with collagens were removed from the analysis. The networks layout was set to force-directed layout.

### Trajectory analysis

Trajectory analysis was performed using Monocle version 2.2.0 ^23^. We ordered the cells onto a pseudotime trajectory based on the union of highly variable genes obtained from all cells, as well as those from only healthy or asthmatic donors.

### Supervised analyses using GWAS genes

Asthma-associated GWAS gene list was collected using the GWAS Catalog of EMBL-EBI searching for the term asthma (https://www.ebi.ac.uk/gwas/). The list was downloaded on 8th of February 2018. We took the genes that are in the top 50 hits of our single-cell DE marker list (either epithelial or non-epithelial) and asthma-associated GWAS list (the “matched” gene list). We then hierarchically clustered the expression matrix of the matched gene list along its rows (genes) and columns (single cells) and represented this as a heatmap.

### Neuroendocrine cell identification

Neuroendocrine cells were identified by the expression of CHGA. Any cell expressing any amount of CHGA was classified as a neuroendocrine cell.

### OMIM search for lung diseases

We searched the clinical synopses with known molecular basis in the Online Mendelian Inheritance in Man (OMIM) database^®^ for the following terms: ‘pulm*’ or ‘bronchi*’ or ‘alveol*’ or ‘surfactant’ and retrieved 337 entries. These terms were chosen to minimise the return of genetic conditions causing respiratory insufficiency as a consequence of neuromuscular dysfunction, skeletal dysplasia (small rib cage) or lung segmentation defects arising in early embryogenesis. These 337 entries were then manually curated to identify those conditions with features affecting the bronchial tree, alveoli, lung parenchyma and pulmonary vasculature. On manual review, entries containing terms such as ‘alveolar ridge’ of the jaw and ‘pulmonary valve stenosis’ and ‘pulmonary embolism’, but no terms related to primary pulmonary disorders, were excluded from further consideration. Syndromes caused by chromosomal disorder or contiguous gene deletion were excluded.

### Statistical methods

For 10x samples comparing healthy versus asthma, we used Fisher’s exact test corrected for multiple testing with Bonferroni method. Normalised CD4 cluster proportions were analysed via paired t-tests corrected for multiple testing with Holm-Sidak method.

## References

1. Bousquet, J., Dahl, R. & Khaltaev, N. Global Alliance against Chronic Respiratory Diseases. Eur. Respir. J. 29, 233–239 (2007).

2. Regev, A. et al. The Human Cell Atlas. Elife 6, (2017).

3. Franks, T. J. et al. Resident cellular components of the human lung: current knowledge and goals for research on cell phenotyping and function. Proc. Am. Thorac. Soc. 5, 763–766 (2008).

4. Montoro, D. T. et al. A revised airway epithelial hierarchy includes CFTR-expressing Nature 560, 319–324 (2018).

5. Plasschaert, L. W. et al. A single-cell atlas of the airway epithelium reveals the CFTR-rich pulmonary ionocyte. Nature 560, 377–381 (2018).

6. Tata, P. R. & Rajagopal, J. Plasticity in the lung: making and breaking cell identity. Development 144, 755–766 (2017).

7. Macosko, E. Z. et al. Highly Parallel Genome-wide Expression Profiling of Individual Cells Using Nanoliter Droplets. Cel l 161, 1202–1214 (2015).

8. Mereu, E. et al. matchSCore: Matching Single-Cell Phenotypes Across Tools and Experiments. bioRxiv 314831 (2018). doi:http://dx.doi.org/10.1101/314831” 10.1101/314831

9. Bisset, L. R. & Schmid-Grendelmeier, P. Chemokines and their receptors in the pat of allergic asthma: progress and perspective. Curr. Opin. Pulm. Med. http://paperpile.com/b/zspAc8/6h1R” 11, p35–42 (2005).

10. Colvin, R. A. et al. Synaptotagmin-mediated vesicle fusion regulates cell migration. Nat. Immunol. 11, 495–502 (2010).

11. Urawa, M. et al. Protein S is protective in pulmonary fibrosis. J. Thromb. Haemost. h 14, 1588–1599 (2016).

12. Wujak, L. A. http://paperpile.com/b/zspAc8/kqgK” et al. FXYD1 negatively regulates Na(+)/K(+)-ATPase activity in lung alveolar epithelial cells. Respir. Physiol. Neurobiol. 220, 54–61 (2016).

13. Krotova, K. et al. Alpha-1 Antitrypsin-Deficient Macrophages Have Increased Matriptase-Mediated Proteolytic Activity. Am. J. Respir. Cell Mol. Biol. 57, 238–247 (2017).

14. Vogl, T. et al. S100A12 is expressed exclusively by granulocytes and acts independently from MRP8 and MRP14. J. Biol. Chem. 274, 25291–25296 (1999).

15. Mitchell, A. et al. LILRA5 is expressed by synovial tissue macrophages in rheumatoid arthritis, selectively induces pro-inflammatory cytokines and IL-10 and is regulated by TNF-alpha, IL-10 and IFN-gamma. Eur. J. Immunol. 38, 3459–3473 (2008).

16. Condon, T. V., Sawyer, R. T., Fenton, M. J. & Riches, D. W. H. Lung dendritic cells at the innate-adaptive immune interface. J. Leukoc. Biol. 90, 883–895 (2011).

17. Holgate, S. T. et al. Asthma. Nat Rev Dis Primers 1, 15025 (2015).

18. Lopez-Guisa, J. M. et al. Airway epithelial cells from asthmatic children differentially express proremodeling factors. J. Allergy Clin. Immunol. 129, 990–7.e6 (2012).

19. Alcala, S. E. et al. Mitotic asynchrony induces transforming growth factor-β1 secretion from airway epithelium. Am. J. Respir. Cell Mol. Biol. 51, 363–369 (2014).

20. Harkness, L. M., Ashton, A. W. & Burgess, J. K. Asthma is not only an airway disease, but also a vascular disease. Pharmacol. Ther. 148, 17–33 (2015).

21. Balzar, S. et al. Mast cell phenotype, location, and activation in severe asthma. Data from the Severe Asthma Research Program. Am. J. Respir. Crit. Care Med. 183, 299–309 (2011).

22. Truyen, E. et al. Evaluation of airway inflammation by quantitative Th1/Th2 cytokine mRNA measurement in sputum of asthma patients. Thorax 61, 202–208 (2006).

23. Trapnell, C. et al. The dynamics and regulators of cell fate decisions are revealed by pseudotemporal ordering of single cells. Nat. Biotechnol. 32, 381–386 (2014).

24. Erle, D. J. & Sheppard, D. The cell biology of asthma. J. Cell Biol. 205, 621–631 (2014).

25. Danahay, H. et al. Notch2 is required for inflammatory cytokine-driven goblet cell metaplasia in the lung. Cell Rep. 10, 239–252 (2015).

26. Gomi, K., Arbelaez, V., Crystal, R. G. & Walters, M. S. Activation of NOTCH1 or NOTCH3 signalling skews human airway basal cell differentiation toward a secretory pathway. PLoS One 10, e0116507 (2015).

27. Ordovas-Montanes, J. et al. Allergic inflammatory memory in human respiratory epithelial progenitor cells. Nature 560, 649–654 (2018).

28. Luo, W. et al. Airway Epithelial Expression Quantitative Trait Loci Reveal Genes Underlying Asthma and Other Airway Diseases. Am. J. Respir. Cell Mol. Biol. 54, 177–187 (2016).

29. Wu, C. A. http://paperpile.com/b/zspAc8/lm0i” et al. Bronchial epithelial cells produce IL-5: implications for local immune responses in the airways. Cell. Immunol. 264, 32–41 (2010).

30. Laitinen, L. A., Laitinen, A. & Haahtela, T. Airway mucosal inflammation even in patients with newly diagnosed asthma. Am. Rev. Respir. Dis. 147, 697–704 (1993).

31. Arima, M. & Fukuda, T. Prostaglandin D2and TH2 Inflammation in the Pathogenesis of Bronchial Asthma. Korean J. Intern. Med. 26, 8 (2011).

32. Xue, L. et al. Prostaglandin D2 activates group 2 innate lymphoid cells through chemoattractant receptor-homologous molecule expressed on TH2 cells. J. Allergy Clin. Immunol. 133, 1184–1194 (2014).

33. Dougherty, R. H. et al. Accumulation of intraepithelial mast cells with a unique protease phenotype in T(H)2-high asthma. J. Allergy Clin. Immunol. 125, 1046–1053.e8 (2010).

34. Hol, B. E., van de Graaf, E. A., Out, T. A., Hische, E. A. & Jansen, H. M. IgM in the airways of asthma patients. Int. Arch. Allergy Appl. Immunol. 96, 12–18 (1991).

35. Peebles, R. S., Jr, Liu, M. C., Lichtenstein, L. M. & Hamilton, R. G. IgA, IgG and IgM quantification in bronchoalveolar lavage fluids from allergic rhinitics, allergic asthmatics, and normal subjects by monoclonal antibody-based immunoenzymetric assays. J. Immunol. Methods 179, 77–86 (1995).

36. Muehling, L. M., Lawrence, M. G. & Woodfolk, J. A. Pathogenic CD4+ T cells in patients with asthma. J. Allergy Clin. Immunol. 140, 1523–1540 (2017).

37. Oja, A. E. et al. Trigger-happy resident memory CD4+ T cells inhabit the human lungs. Mucosal Immunol. 11, 654–667 (2018).

38. Kumar, B. V. et al. Human Tissue-Resident Memory T Cells Are Defined by Core Transcriptional and Functional Signatures in Lymphoid and Mucosal Sites. Cell Rep. 20, 2921–2934 (2017).

39. Mitson-Salazar, A. et al. Hematopoietic prostaglandin D synthase defines a proeosinophilic pathogenic effector human T(H)2 cell subpopulation with enhanced function. J. Allergy Clin. Immunol. 137, 907–18.e9 (2016).

40. Wambre, E. et al. A phenotypically and functionally distinct human TH2 cell subpopulation is associated with allergic disorders. Sci. Transl. Med. 9, (2017).

41. Lam, E. P. S. et al. IL-25/IL-33-responsive TH2 cells characterize nasal polyps with a default TH17 signature in nasal mucosa. J. Allergy Clin. Immunol. 137, 1514–1524 (2016).

42. Vento-Tormo, R. et al. Single-cell reconstruction of the early maternal–fetal interface in humans. Nature 563, 347–353 (2018).

43. Weckmann, M., Kopp, M. V., Heinzmann, A. & Mattes, J. Haplotypes covering the TNFSF10 gene are associated with bronchial asthma. Pediatr. Allergy Immunol. 22, 25–30 (2011).

44. Harada, M. et al. Thymic stromal lymphopoietin gene promoter polymorphisms are associated with susceptibility to bronchial asthma. Am. J. Respir. Cell Mol. Biol. 44, 787–793 (2011).

45. Grotenboer, N. S., Ketelaar, M. E., Koppelman, G. H. & Nawijn, M. C. Decoding asthma: translating genetic variation in IL33 and IL1RL1 into disease pathophysiology. J. Allergy Clin. Immunol. 131, 856–865 (2013).

46. Gerovac, B. J. & Fregien, N. L. IL-13 Inhibits Multicilin Expression and Ciliogenesis via Janus Kinase/Signal Transducer and Activator of Transcription Independently of Notch Cleavage. Am. J. Respir. Cell Mol. Biol. 54, 554–561 (2016).

47. Holgate, S. T. et al. Epithelial-mesenchymal communication in the pathogenesis of chronic asthma. Proc. Am. Thorac. Soc. 1, 93–98 (2004).

48. Butler, A., Hoffman, P., Smibert, P., Papalexi, E. & Satija, R. Integrating single-cell transcriptomic data across different conditions, technologies, and species. Nat. Biotechnol. 36, 411–420 (2018).

49. Song, J. et al. Aberrant DNA methylation and expression of SPDEF and FOXA2 in airway epithelium of patients with COPD. Clin. Epigenetics 9, 42 (2017).

50. Heijink, I. H. et al. Down-regulation of E-cadherin in human bronchial epithelial cells leads to epidermal growth factor receptor-dependent Th2 cell-promoting activity. J. Immunol. 178, 7678–7685 (2007).

51. Picelli, S. et al. Full-length RNA-seq from single cells using Smart-seq2. Nat. Protoc. 9, 171– 181 (2014).

52. Wu, T. D. & Nacu, S. Fast and SNP-tolerant detection of complex variants and splicing in short reads. Bioinformatics 26, 873–881 (2010).

53. Dobin, A. et al. STAR: ultrafast universal RNA-seq aligner. Bioinformatics 29, 15–21 (2013).

54. van den Brink, S. C.; et al. Single-cell sequencing reveals dissociation-induced gene; expression in tissue subpopulations. Nat. Methods 14, 935–936 (2017).

55. Young, M. D. & Behjati, S. SoupX removes ambient RNA contamination from droplet based single cell RNA sequencing data. bioRxiv (2018).

