## Supplementary_figures for "A cellular census of healthy lung and asthmatic airway wall identifies novel cell states in health and disease"

### Vieira Braga et al 2019

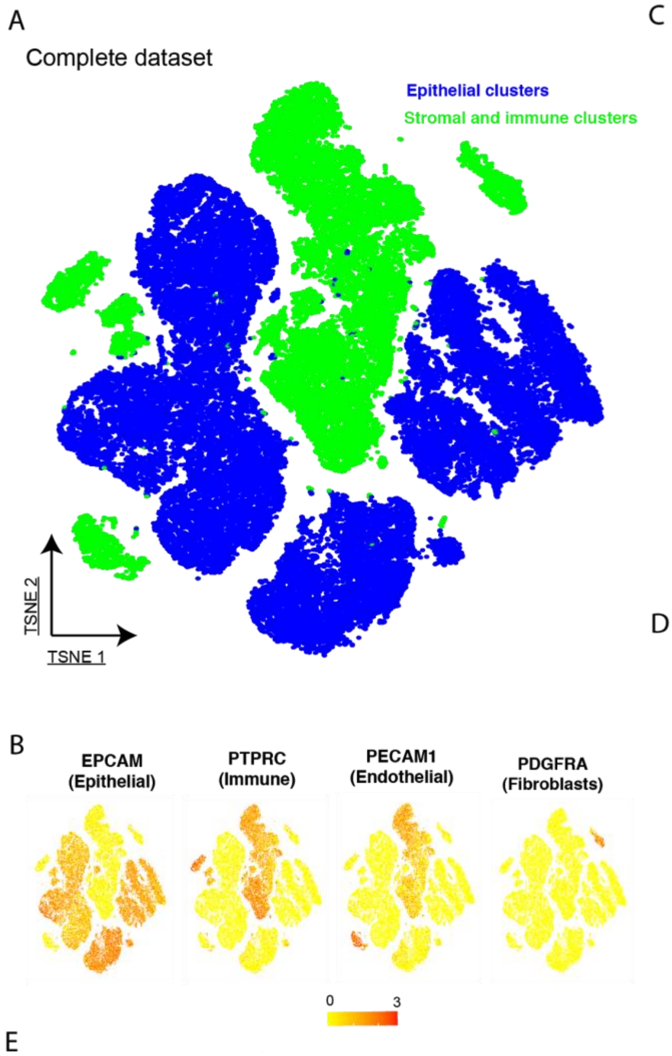


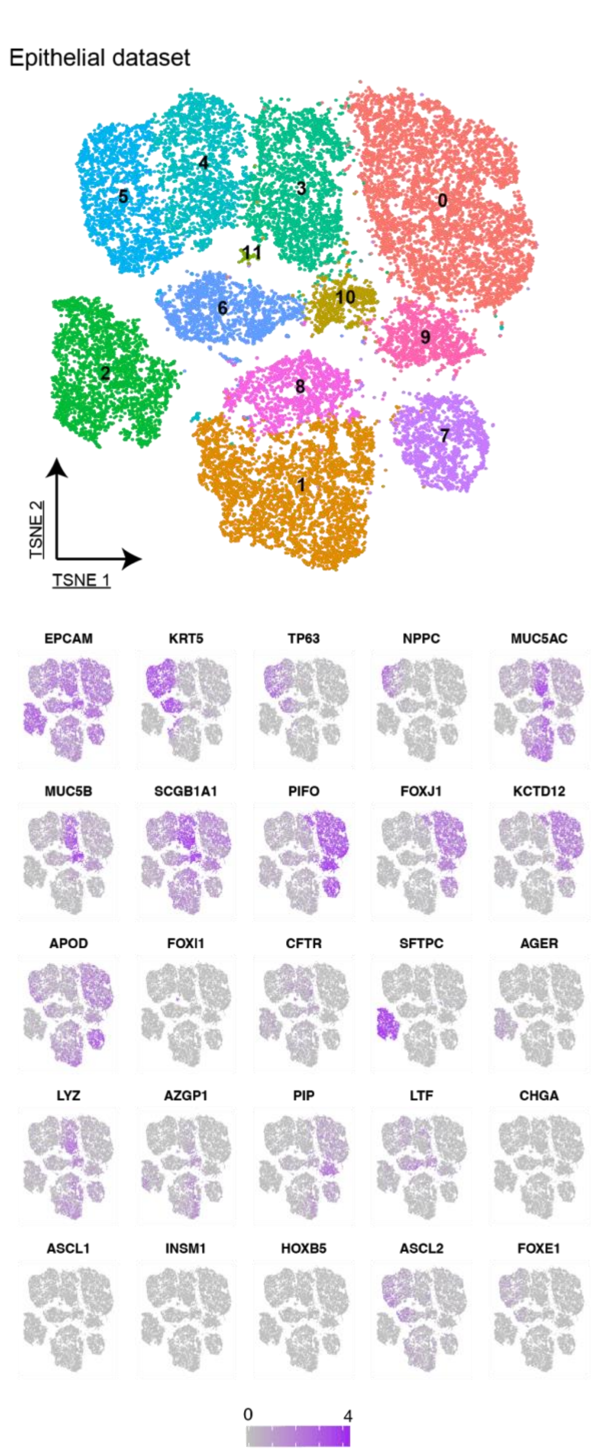


E

| Cluster Number | Cell type assigned | Positive markers | Negative Markers |
|---|---|---|---|
| 0<br>9 | **Ciliated 1** | FOXJ1, PIFO, KCTD12 high | APOD low |
| 2 | **Type 2 alveolar** | SFTPC | |
| 3<br>10 | **Club** | SCGB1A1 high, MUC5AC high, MUC5B high | |
| 5 | **Basal 1** | KRT5, TP63 high, NPPC high | |
| 6<br>4 | **Basal 2** | KRT5, TP63 low | NPPC |
| 1 | **Goblet 1** | MUC5AC high | MUC5B low, SCGB1A1 low |
| 8 | **Goblet 2** | MUC5AC high, CSF3 high | MUC5B low, SCGB1A1 low |
| 7 | **Ciliated 2** | APOD high | |
| 11 | **Ionocytes** | FOXI1, CFTR | |

**Extended Data Figure 1 – Unbiased clustering of upper airway, lower airway and parenchymal lung tissue cells and strategy to divide the dataset used in figures 1 and 2. (A)** tSNEplot depicting unbiased cluster assignment of combined dataset of parenchyma and upper and lower airways, highlighting EPCAM-high cell clusters (epithelial cells) in blue and other cell types in green. **(B)** TSNE as in (A), highlighting expression of cell lineage markers. **(C)** TSNE plot of only epithelial cells , as defined in (A). **(D)** TSNE as in C, showing expression of individual genes used for cell type assignment of different epithelial subsets. **(E)** Table depicting cell type assignment of clusters identified in (C). Alveolar type 1 cells were not identified in an individual cluster in this analysis and they were selected by the high expression of the type 1 marker AGER.

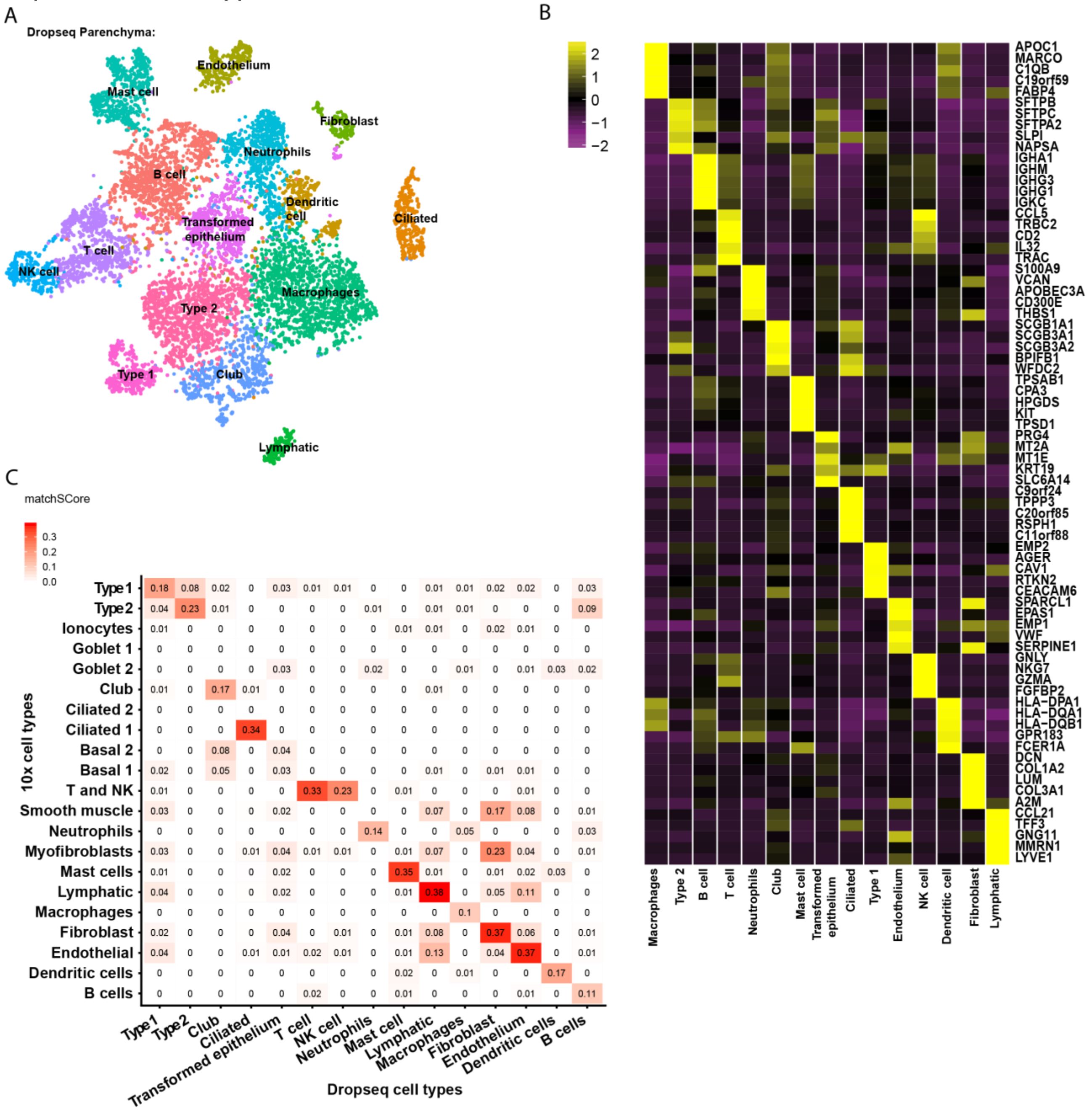


**Extended Data Figure 2**. **Resection lung material analysis via Dropseq (A)** tSNE showing single cells obtained from lung resection samples analysed using Dropseq. **(B)** Heatmap depicting top differentially expressed genes by log fold change among the clusters present in (A). **(C)** Matchscore analysis comparing the clusters present in the Dropseq analysis of lung resection material to the clusters identified in figures 1 and 2.

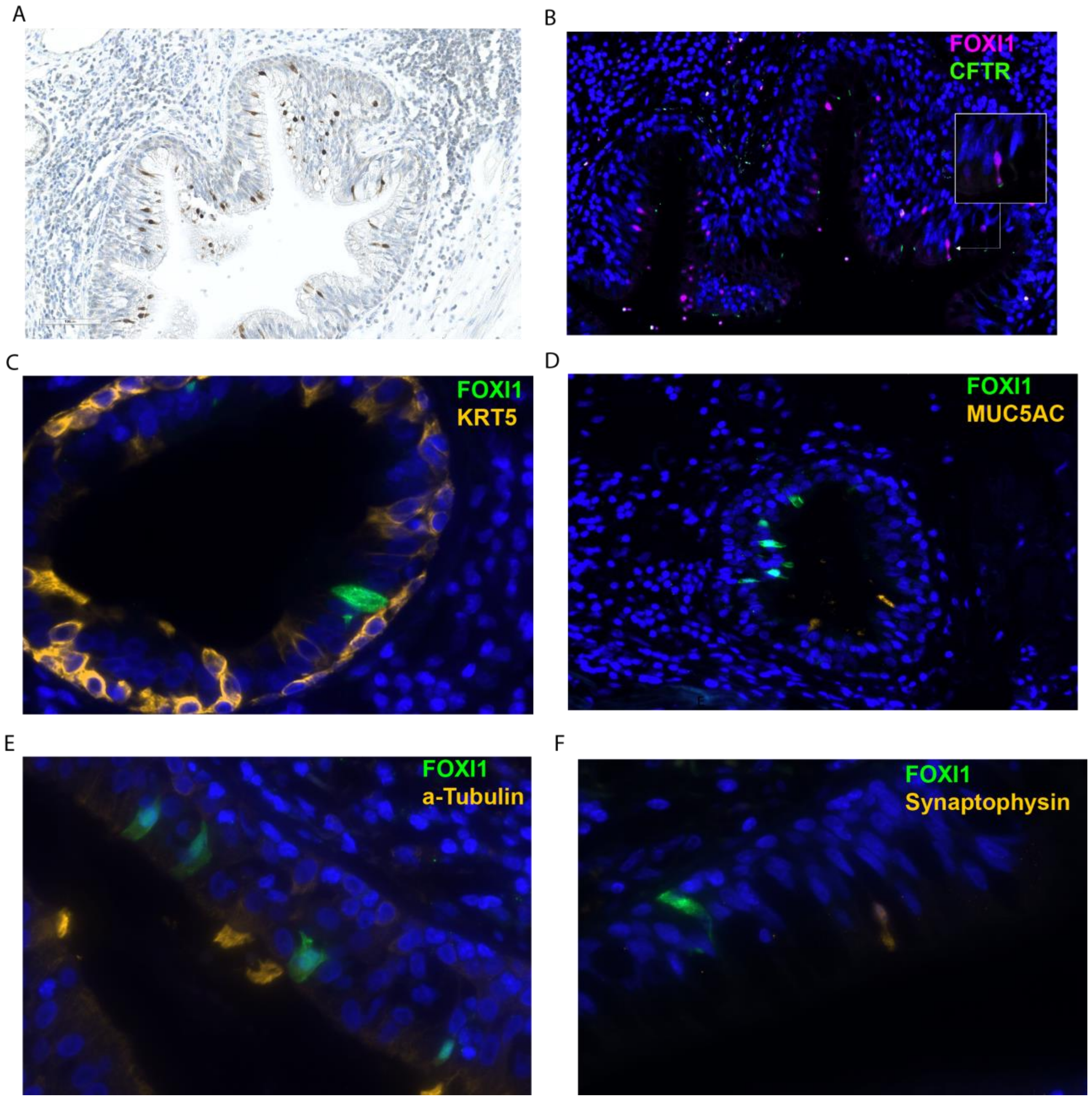


**Extended Data Figure 3. Ionocyte cell identity and molecular profile** (**A**) Immunohistochemistry staining for FOXI1, using haematoxylin as counterstain. **(B/C)** Fluorescent staining for ionocyte markers: mouse anti-FOXI1 and rabbit anti-CFTR **(B)** and mouse anti-FOXI1 and rabbit anti-ATP6V1G3 **(C)** D-G) Fluorescent staining for FOXI1 (ionocyte-specific) and other epithelial markers: **(D)** neuroendocrine cell marker (rabbit anti-synaptophysin), **(E)** ciliated cell marker (mouse anti-α-Tubulin) (F) basal cell marker (mouse anti-α-KRT5) **(G)** secretory cell marker (mouse anti-MUC5AC) DAPI (blue) stains nuclei.

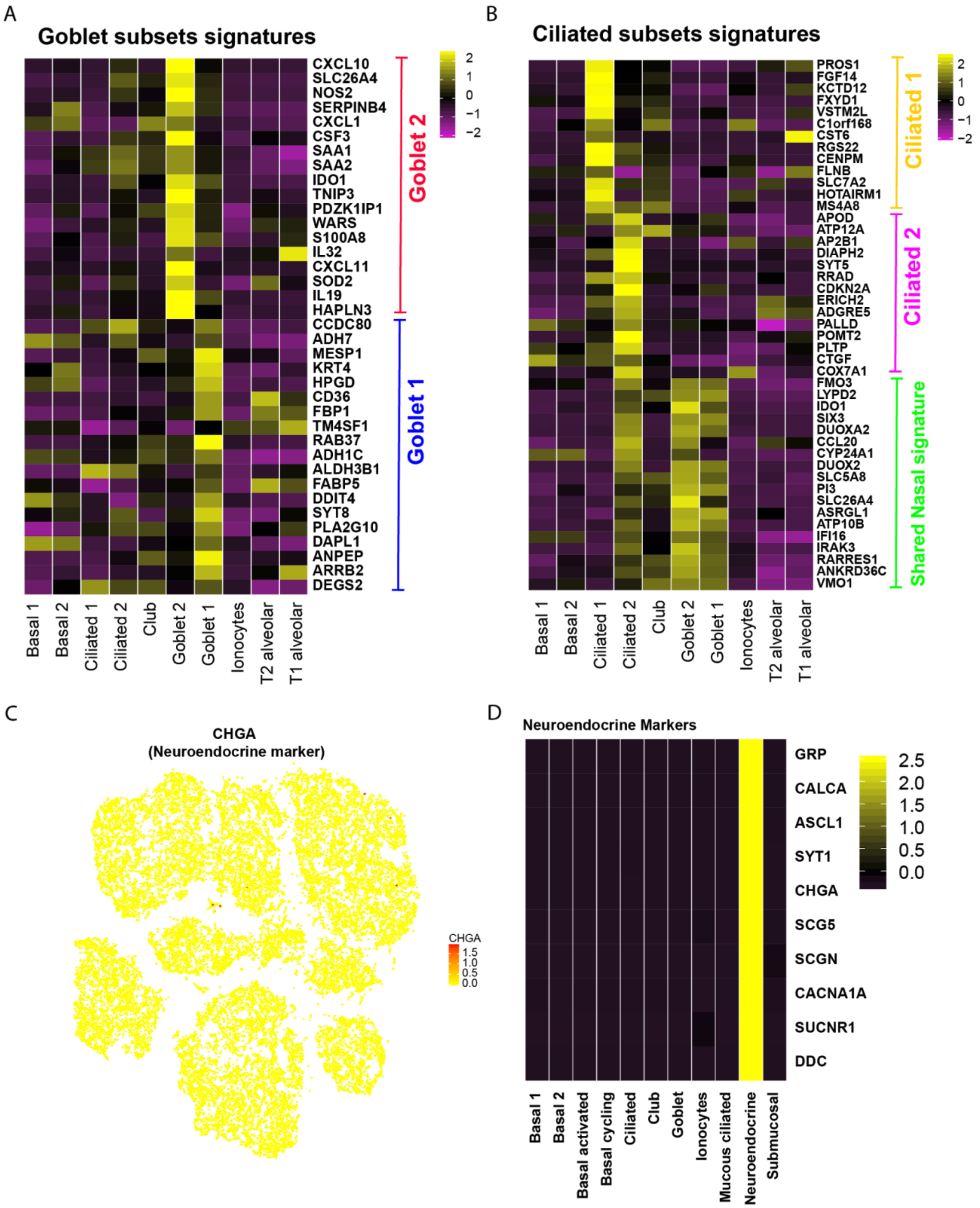


**Extended Data Figure 4. Marker gene analysis of specific goblet, ciliated and neuroendocrine cells in human lungs. (A)** Heatmap depicting the expression of marker genes identified by differential expression analysis comparing Goblet 1 *versus* Goblet 2 cell types. **(B)** Heatmap depicting the expression of marker genes identified by differential expression analysis. Ciliated 1 genes and shared nasal signature genes generated by comparing Ciliated 1 *versus* Ciliated 2. Ciliated 2 genes generated by comparing Ciliated 2 *versus* the combination of Ciliated 1, Goblet 1 and Goblet 2, in order to subtract the nasal signature. **(C)** tSNE plot depicting the expression of the neuroendocrine marker CHGA. **(D)** Heatmap depicting the expression of Neuroendocrine markers in neuroendocrine cells as well as in the other clusters identified in our analysis. Neuroendocrine cells identified based on CHGA expression. Neuroendocrine markers obtained from Ordovas-Montanes *et al.* *Nature* **560,** 2018.

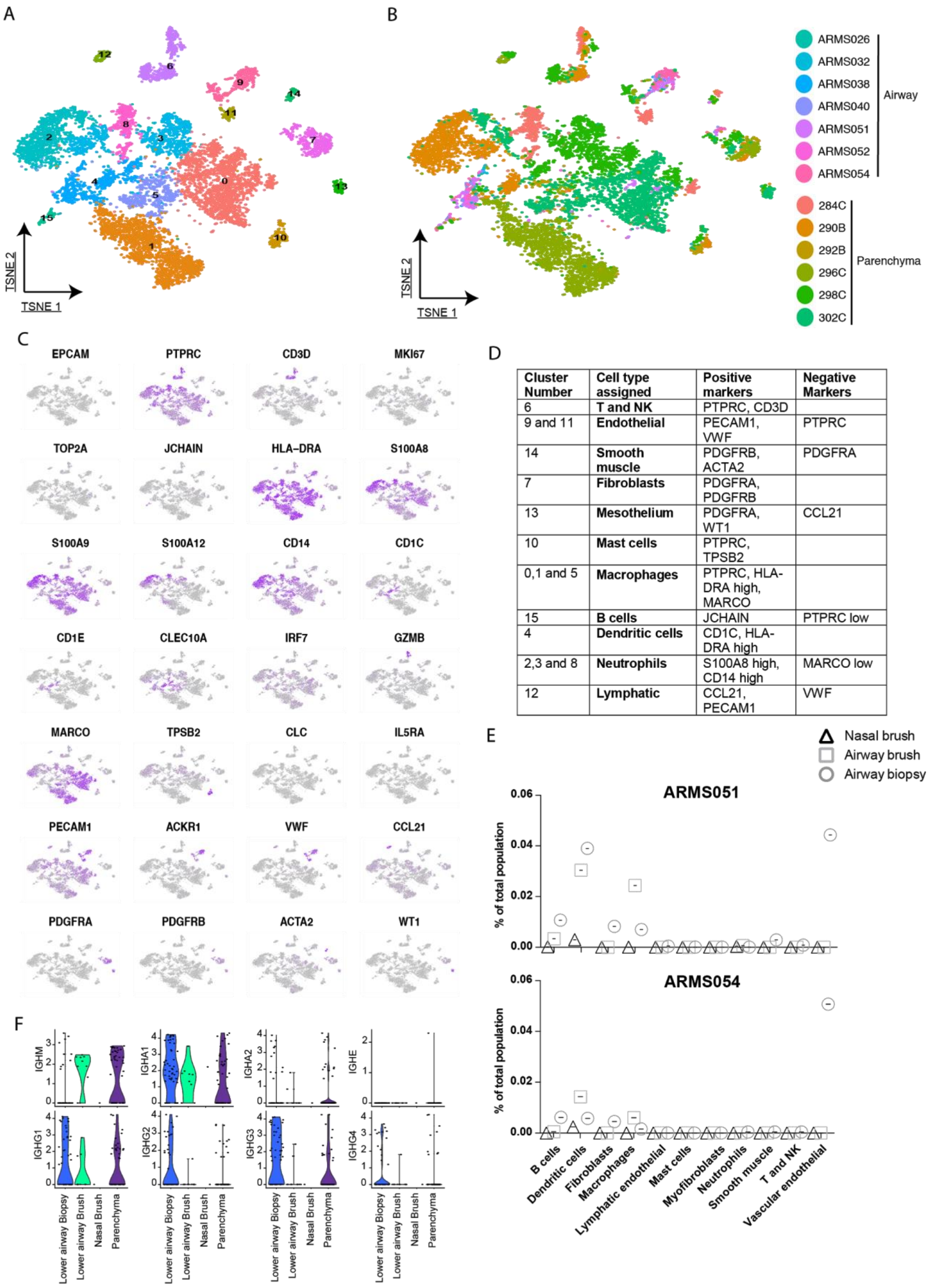


| Cluster Number | Cell type assigned | Positive markers | Negative Markers |
|---|---|---|---|
| 6 | **T and NK** | PTPRC, CD3D | |
| 9 and 11 | **Endothelial** | PECAM1, VWF | PTPRC |
| 14 | **Smooth muscle** | PDGFRB, ACTA2 | PDGFRA |
| 7 | **Fibroblasts** | PDGFRA, PDGFRB | |
| 13 | **Mesothelium** | PDGFRA, WT1 | CCL21 |
| 10 | **Mast cells** | PTPRC, TPSB2 | |
| 0,1 and 5 | **Macrophages** | PTPRC, HLA-DRA high, MARCO | |
| 15 | **B cells** | JCHAIN | PTPRC low |
| 4 | **Dendritic cells** | CD1C, HLA-DRA high | |
| 2,3 and 8 | **Neutrophils** | S100A8 high, CD14 high | MARCO low |
| 12 | **Lymphatic** | CCL21, PECAM1 | VWF |

**Extended Data Figure 5**. **Cell type assignment strategy for the assignment of non-epithelial cells discussed in Figure 2.** (**A)** tSNE coloured by unbiased clustering of the non-epithelial dataset described in Figure 2. **(B)** tSNE depicting single cells coloured by their respective sample origin. **(C)** tSNE as in (A), showing the expression of lineage markers used for cell type assignment. **(D)** Table with the strategy used for cluster cell assignment of the cell types present in figure 2. **(E)** Cluster distribution in the two donors from which we collected paired nasal brushes, airway brushes and airway biopsies. **(F)** Violin plots depicting the expression of immunoglobulin genes in the B cell cluster divided by tissue of origin.

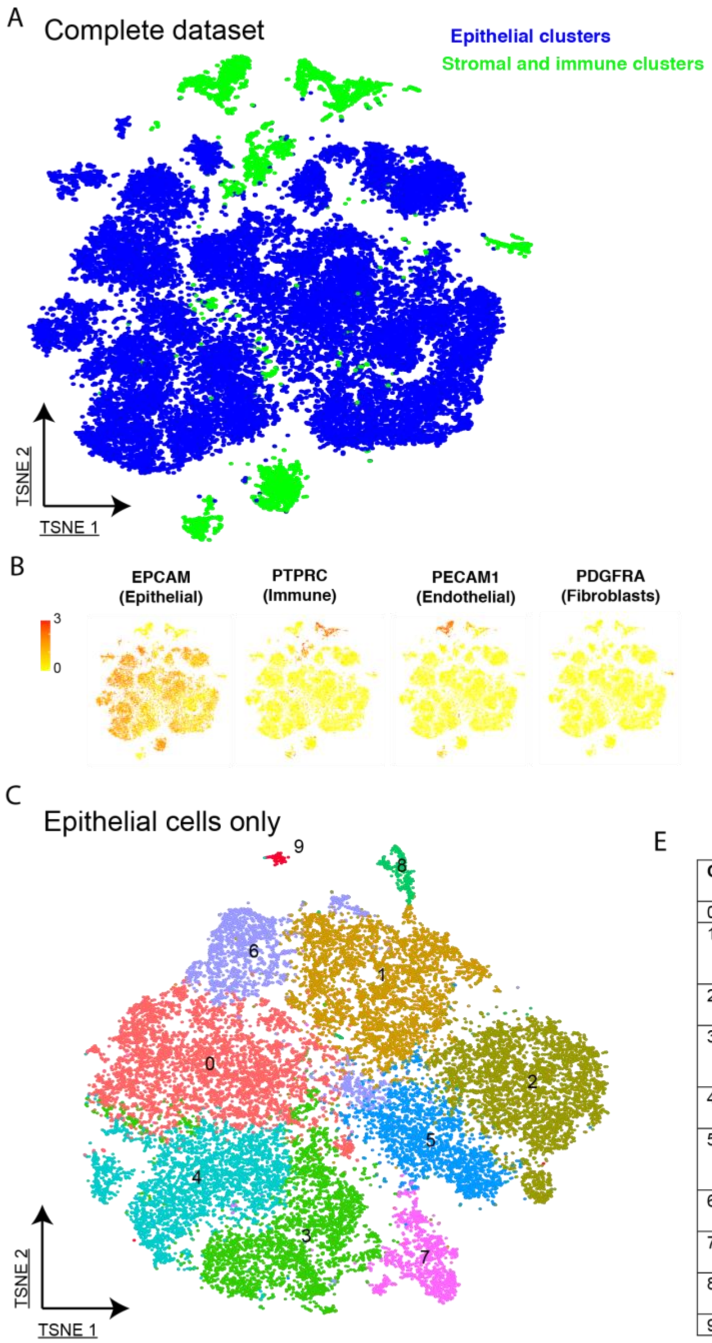


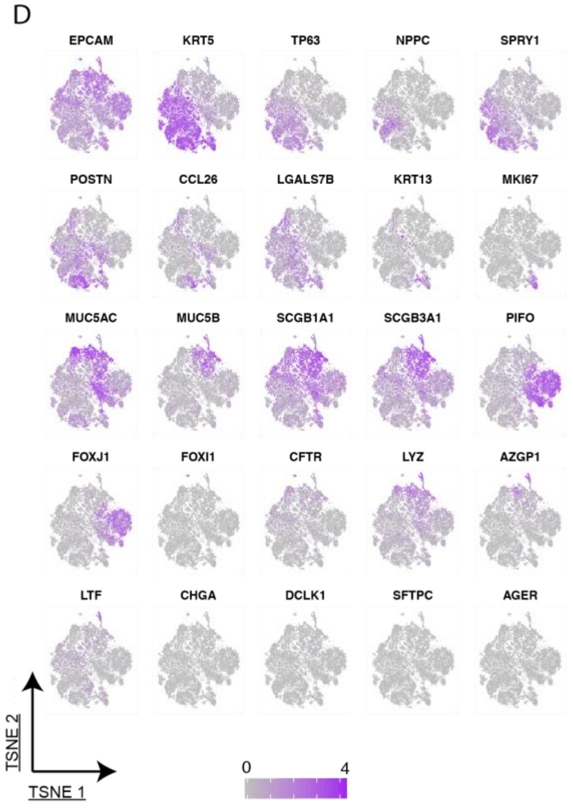


E

| Cluster Number | Cell type assigned | Positive markers | Negative Markers |
|---|---|---|---|
| 0 | **Basal 2** | KRT5, TP63 low | NPPC |
| 1 | **Club** | SCGB1A1 high, MUC5AC high, MUC5B high | |
| 2 | **Ciliated** | PIFO, FOXJ1 | |
| 3 | **Basal activated** | KRT5, TP63 high, SPRY1 high, POSTN high | |
| 4 | **Basal 1** | KRT5, TP63 high, NPPC high | |
| 5 | **Mucous ciliated** | MUC5AC high, PIFO high, FOXJ1 high | MUC5B low, SCGB1A1 low |
| 6 | **Goblet** | MUC5AC high | MUC5B low, SCGB1A1 low |
| 7 | **Basal cycling** | KRT5, TP63 low, MKI67, | |
| 8 | **Submucosal gland** | AZGP1, LYZ,PIP,LTF | |
| 9 | **Ionocytes** | FOXI1, CFTR | |

**Extended Data Figure 6 - Unbiased clustering of lower airway biopsy samples of healthy and asthmatic volunteers and the strategy used to divide the dataset depicted in figures 3 and 4. (A)** tSNEplot depicting unbiased cluster assignment of combined dataset of control and asthma lower airway biopsies, highlighting EPCAM-high cell clusters (epithelial cells) in blue and other cell types in green. **(B)** TSNE as in (A), highlighting expression of cell lineage markers. **(C)** tSNE plot of only epithelial cells , as defined in (A). **(D)** TSNE as in C, showing expression of individual genes used for cell type assignment of different epithelial subsets. **(E)** Table depicting cell type assignment of clusters identified in (C).

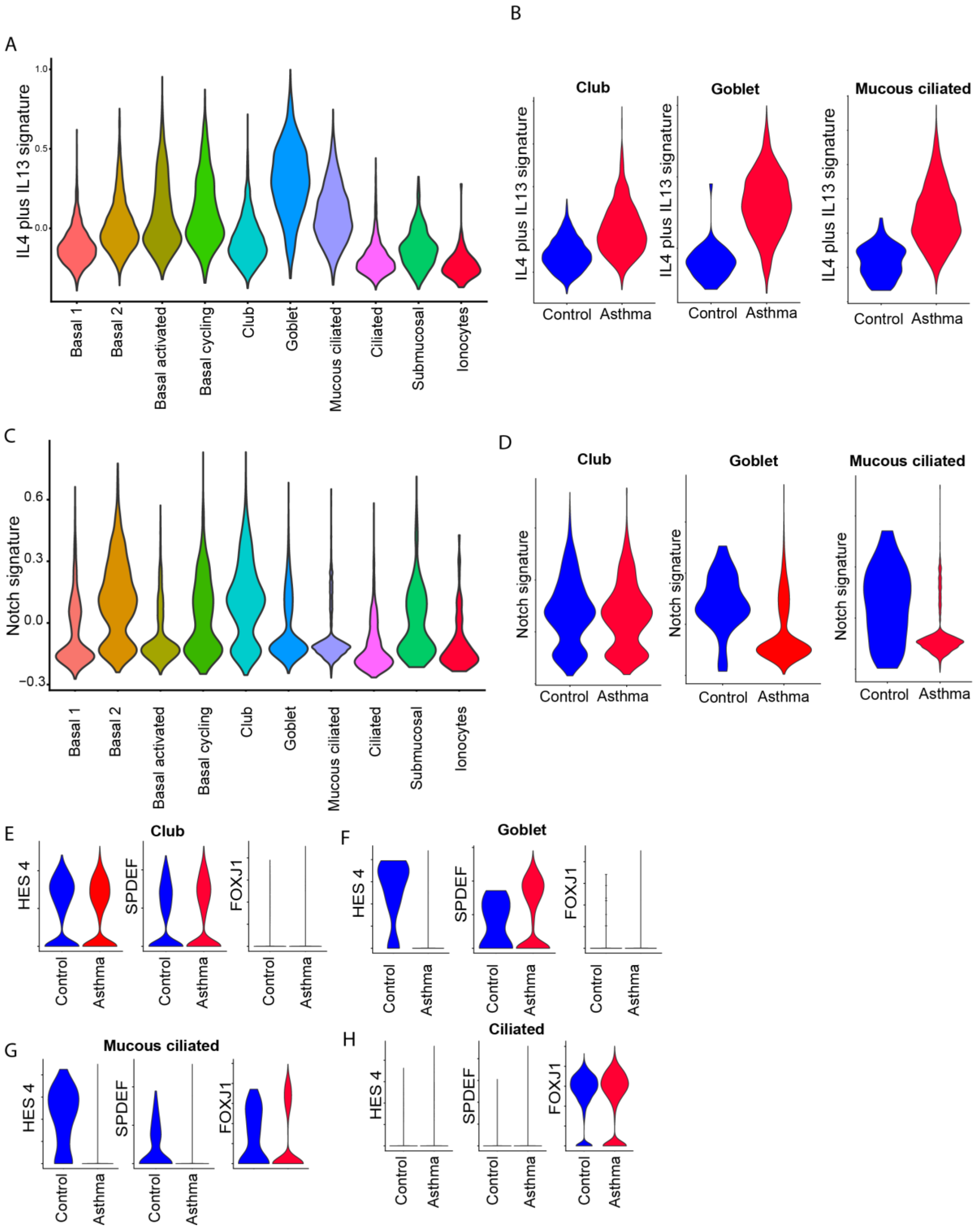


**Extended Data Figure 7. IL4/IL13 and NOTCH driven gene transcription signatures in goblet cell metaplasia in asthma. (A)** Violin plot depicting IL4/IL13 genes signature expression in each cluster of biopsy epithelial cells (Signature obtained from Ordovas-Montanes, et al *Nature* 566**,** 2018). **(B)** Violin plot depicting the expression of the IL4/Il13 signature (as in A) in control and asthma cells of selected clusters. **(C)** Violin plot depicting Notch signature genes (HES4, HES5, HEY1, HEY2, HEYL and NRARP) expression in each cluster of biopsy epithelial cells. **(D)** Violin plot depicting the expression of Notch signature genes (as in C) in control and asthma cells of selected clusters. Expression of secretory specific (HES4 and SPDEF) and ciliated specific (FOXJ1) transcription factors in Club **(E),** Goblet **(F)**, Ciliated **(G)**, and Mucous Ciliated **(H)** cells.

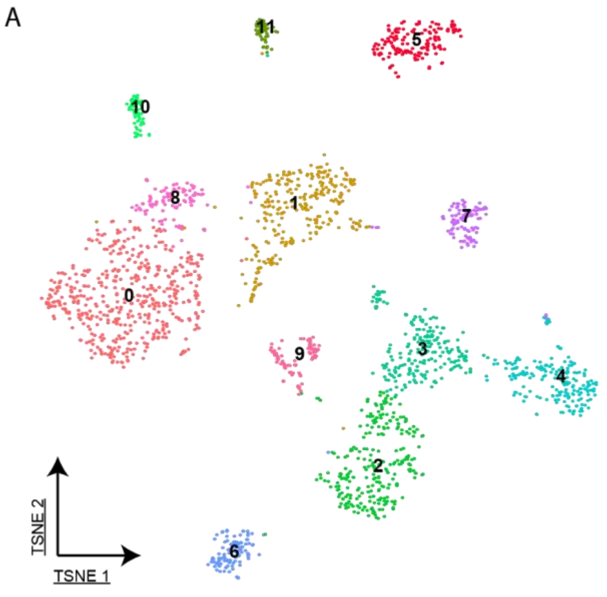


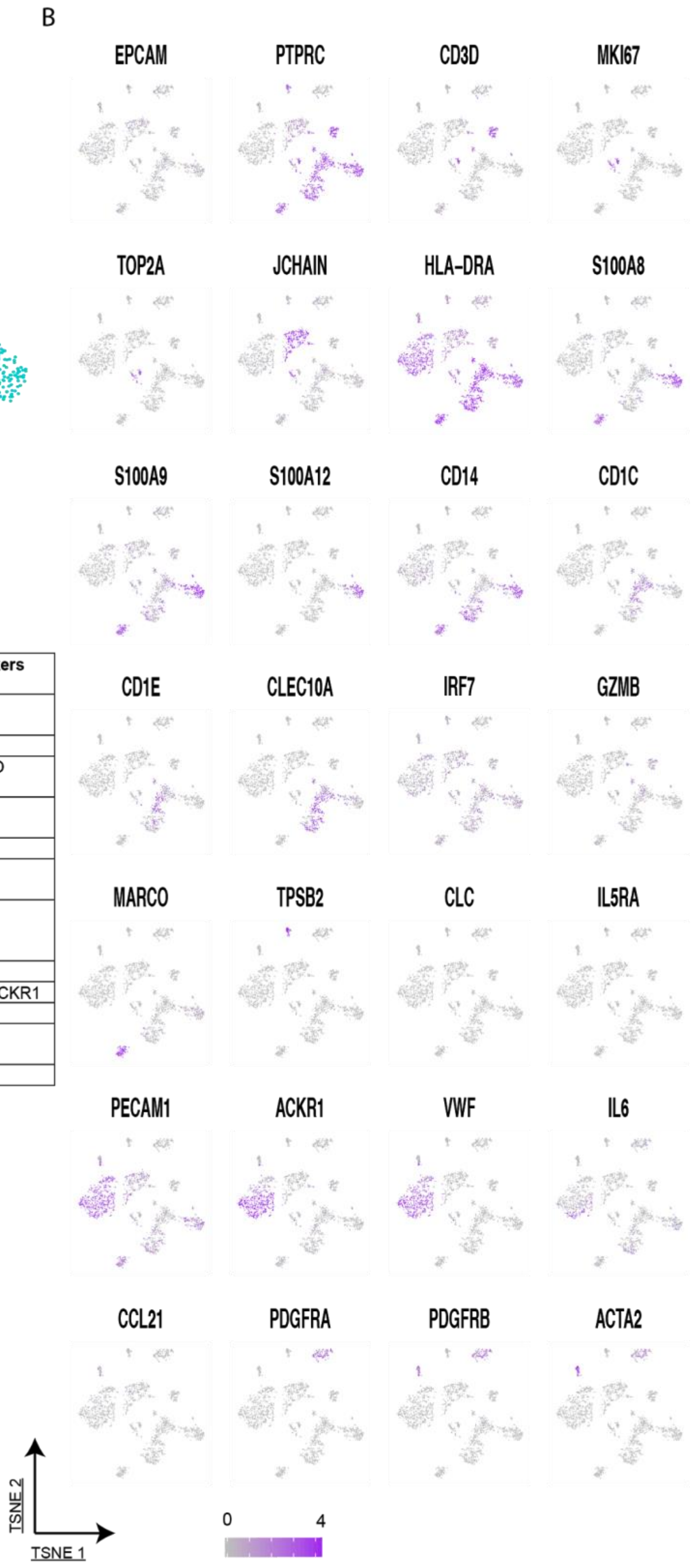


C

| Cluster Number | Cell type assigned | Positive markers | Negative Markers |
|---|---|---|---|
| 0 | **Activated endothelium** | PECAM1, VWF, IL6, ACKR1 | PTPRC |
| 0 | **B cells** | JCHAIN,CD19 | |
| 2 | **Inflammatory DCs** | CLEC10A,CD14, HLA-DRA high | CD1C, MARCO |
| 3 | **DCs** | CLEC10A,CD1C, HLA-DRA high | |
| 4 | **Neutrophils** | MKI67, TOP2A | |
| 5 | **Fibroblasts** | PDGFRA, PDGFRB | |
| 6 | **Luminal Macrophages** | PTPRC, HLA-DRA high, MARCO | |
| 7 | **T cells** | PTPRC, CD3D | |
| 8 | **Endothelium** | PECAM1, VWF | PTPRC, IL6, ACKR1 |
| 9 | **Cycling** | MKI67, TOP2A | |
| 10 | **Smooth muscle** | PDGFRB, ACTA2 | PDGFRA |
| 11 | **Mast cells** | PTPRC, TPSB2 | |

**Extended Data Figure 8. Clustering and cell type assignment of non-epithelial cells in the airways of control and asthma patients.** (**A**) tSNE depicting unbiased clustering of the non-epithelial dataset described in Figure 4. **(B)** tSNE as in (A), showing the expression of lineage markers used for cell type assignment. **(C)** Table with the strategy used for cluster cell assignment of the cell types present in figure 4.

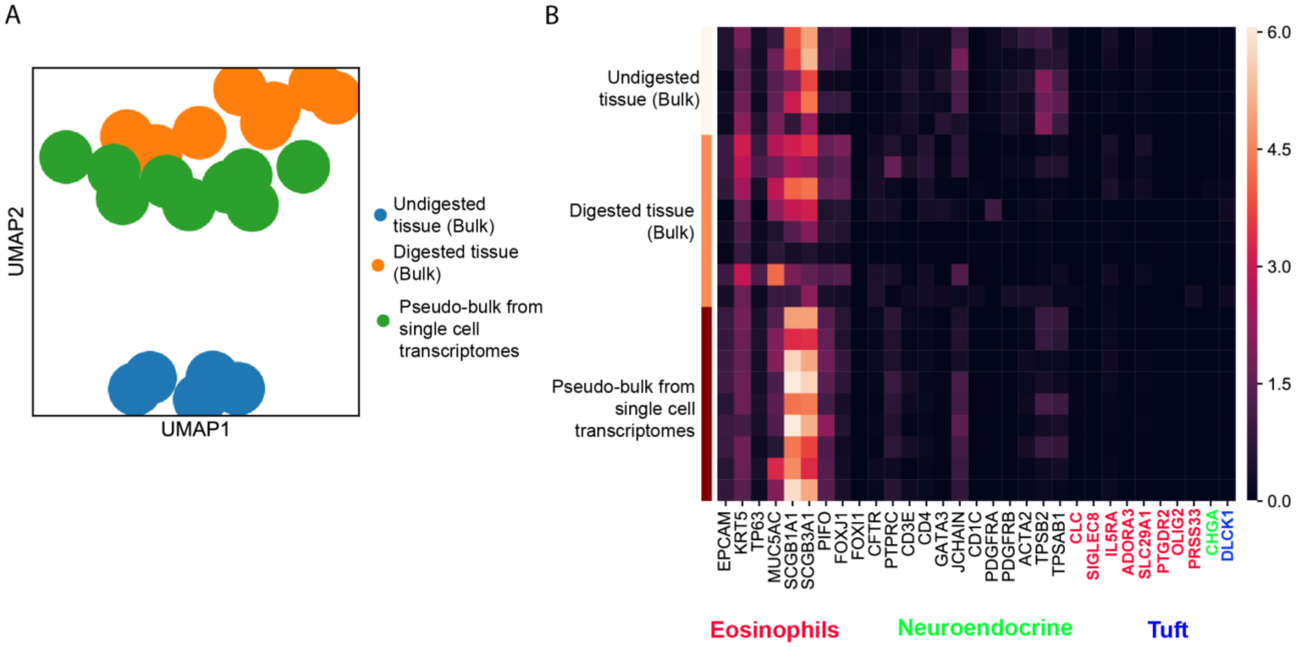


**Extended Data Figure 9. Comparative analysis of bulk transcriptomes versus single-cell RNA seq data on matched airway biopsies. (A)** UMAP displaying bulk transcriptomes of biopsies before digestion (blue), of the single cell suspension obtained from digestion and used to load the 10x microfluidics device (orange) and pseudo-bulk samples generated by collapsing all the data obtained from single cell transcriptomes (green). **(B)** Heatmap depicting expression of several epithelial and non-epithelial cell markers, as shown in Extended Data Figures 6 and 8. Gene markers of rare cells (Neuroendocrine, Eosinophils and Tuft) not identified in our single cell clusters depicted in specific colours.

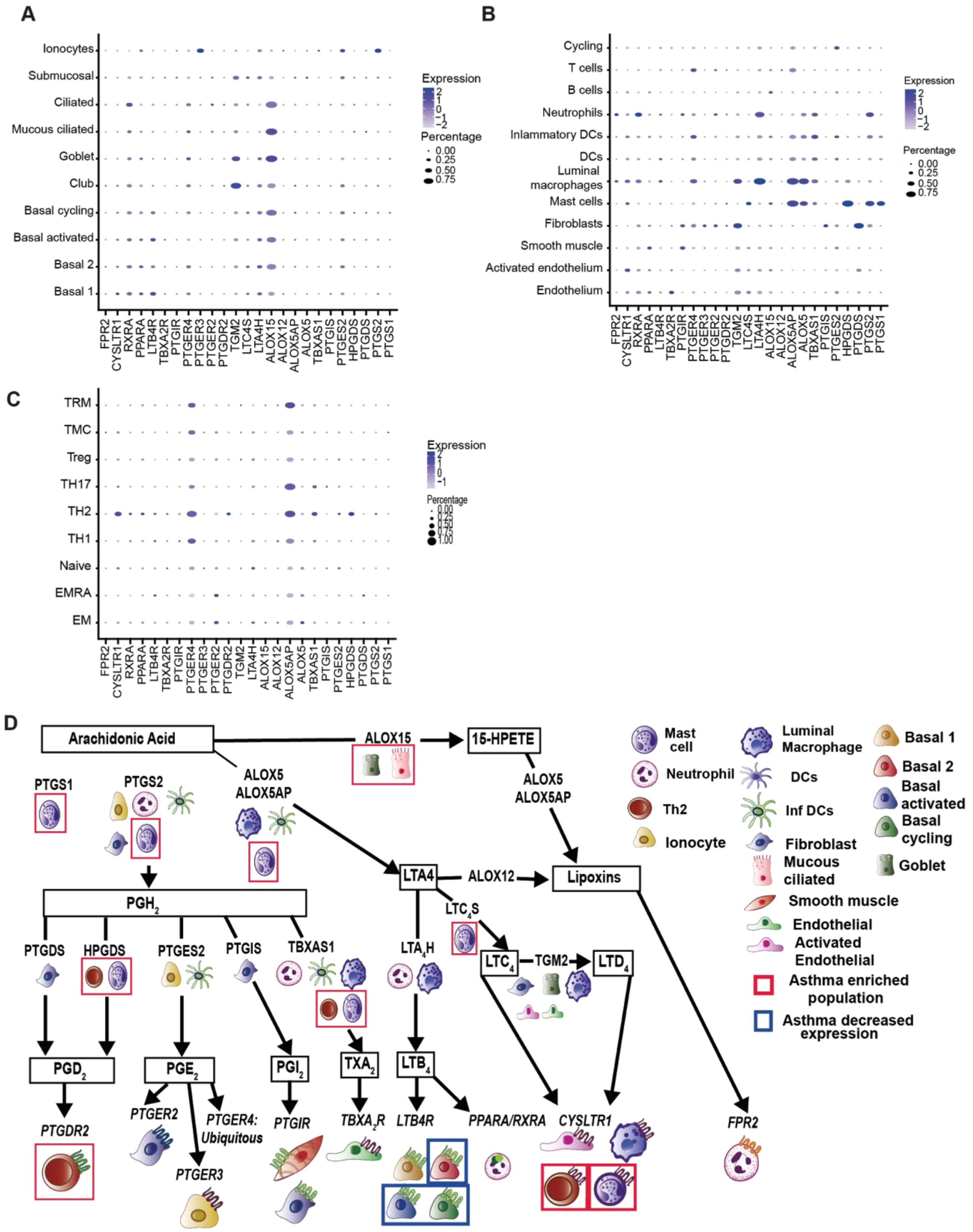


**Extended Data Figure 10. Enzymes of the eicosanoid pathway are unequally distributed among several cell types in asthma.** Dot plots depicting the genes involved in eicosanoids metabolism and their expression in epithelial cells **(A),** non-epithelial cells **(B)** and CD4 T cells **(C). (D)** Cartoon illustrating the eicosanoids pathways and its receptors and in which cells each gene is expressed in the lungs, with special notation for asthma alterations.

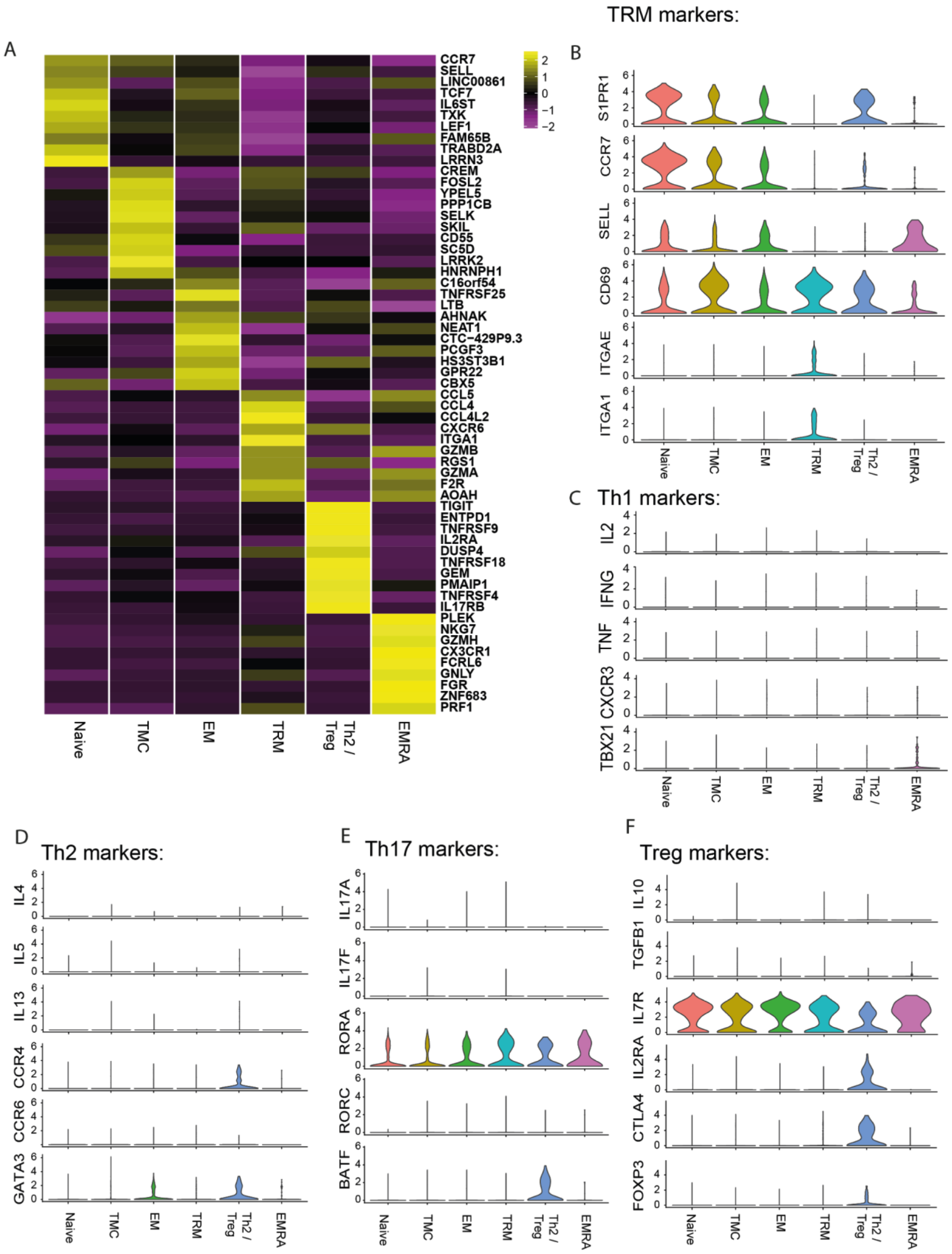


**Extended Data Figure 11. Canonical T cell markers expression in CD4 T cell clusters.** **(A)** Heatmap depicting genes differentially expressed in the five different CD4 T cells clusters identified via unbiased clustering. (**A-F)** Violin plots depicting canonical markers of Tissue Resident Memory Cells **(B)**, Th1 **(C)**, Th2 **(D),** Th17 **(E)** and Treg **(F)** CD4 T cells.

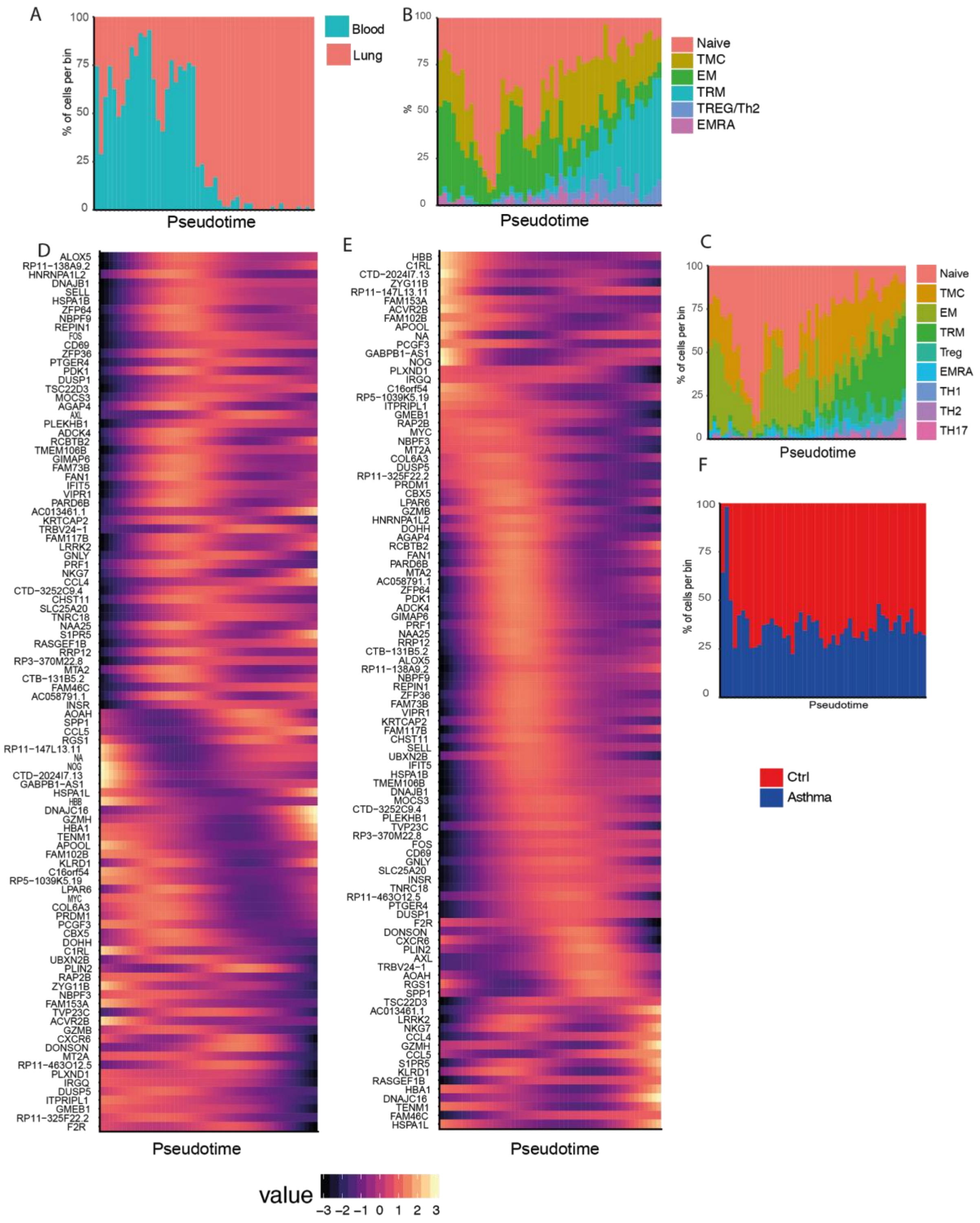


**Extended Data Figure 12. Tissue adaptation signature of CD4 T cells A)** Binned pseudotime analysis depicting proportion of CD4 T cells isolated from blood and lungs. **B)** Cluster cell proportions on the binned pseudotime. **C) and D)** Heatmaps depicting gene expression variations according to pseudotime. **C)** Genes in which the expression decreases as pseudotime progress and **D)** genes with increased expression as pseudotime progresses. **E)** Binned pseudotime showing distribution between asthma and control CD4 T cells.

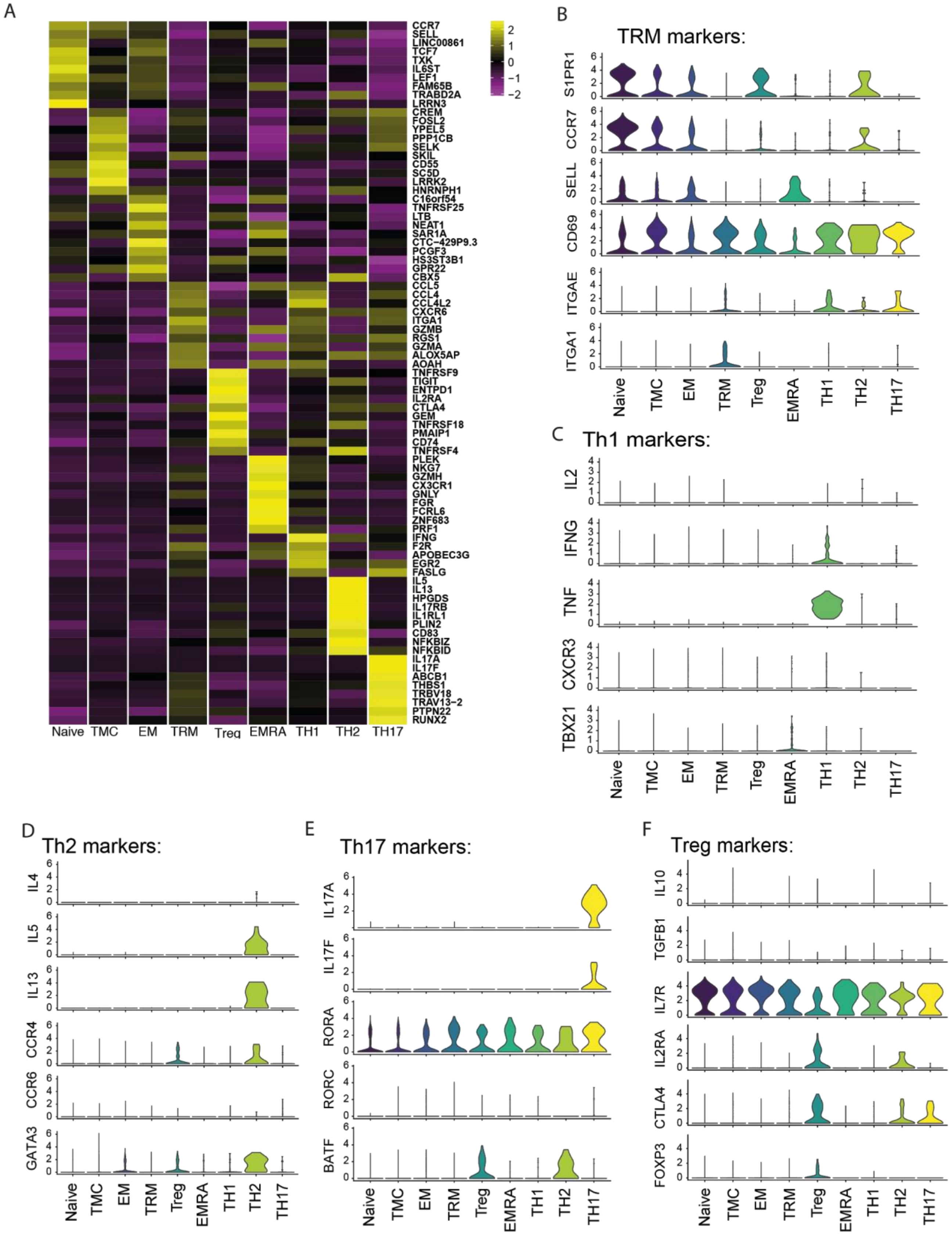




**Extended Data Figure 13. Canonical T cell markers expression in Th CD4 clusters A)** Heatmap depicting genes differentially expressed in the nine clusters of CD4 T cells using an initial unbiased clustering method, followed by manual selection of Th1, Th2 and Th17 clusters based on canonical cytokines. **A-F)** Violin plots depicting canonical markers of Tissue Resident Memory Cells **(B)**, Th1 **(C)**, Th2 **(D),** Th17 **(E)** and Treg **(F)** CD4 T cells.

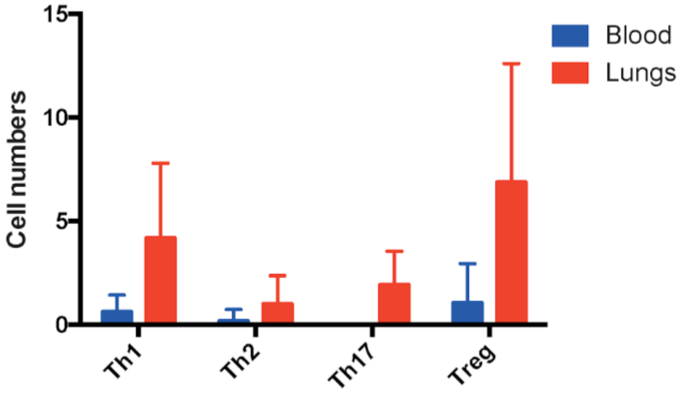


**Extended Data Figure 14. Tissue distribution of classical CD4 T cell subsets. (A)** Bar graph showing the absolute cell numbers of CD4 T cells distributed according to the tissue of origin.
